# Single-cell bisulfite sequencing of spermatozoa from lean and obese humans reveals potential for the transmission of epimutations

**DOI:** 10.1101/2021.07.09.451752

**Authors:** Emil Andersen, Stephen Clark, Lars Ingerslev, Leonidas Lundell, Wolf Reik, Romain Barrès

**Author notes:** To whom correspondence should be addressed: Romain Barrès Panum Institutet 7.7, Blegdamsvej 3B, 2200 Copenhagen N, Denmark.

## Abstract

Epigenetic marks in gametes modulate developmental programming after fertilization. Spermatozoa from obese men exhibit distinct epigenetic signatures compared to lean men, however, whether epigenetic differences are concentrated in a sub-population of spermatozoa or spread across the ejaculate population is unknown. Here, by using whole-genome single-cell bisulfite sequencing on 87 motile spermatozoa from 8 individuals (4 lean and 4 obese), we found that spermatozoa within single ejaculates are highly heterogeneous and contain subsets of spermatozoa with marked imprinting defects. Comparing lean and obese subjects, we discovered methylation differences across two large CpG dense regions located near *PPM1D* and *LINC01237*. These findings confirm that sperm DNA methylation is altered in human obesity and indicate that single ejaculates contain subpopulations of spermatozoa carrying distinct DNA methylation patterns. Distinct epigenetic patterns of spermatozoa within an ejaculate may result in different intergenerational effects and therefore influence strategies aiming to prevent epigenetic-related disorders in the offspring.

## INTRODUCTION

Spermatozoa carry a specific epigenetic signature that controls normal embryonic development (Aston et al., 2015; Hammoud et al., 2009) and abnormalities in the epigenetic pattern of sperm are associated with decreased fertility in humans and raise concern for potential epigenetic transmission to the offspring (Conine et al., 2018; Jenkins et al., 2016; Kaneda et al., 2004; Vasiliauskaitė et al., 2018). Methylation of DNA on the fifth carbon of cytosine (5mC) is an extensively studied epigenetic mark that controls gene expression in time and space, and for which accumulating evidence indicates a role in embryonic development (Peat et al., 2014; Tang et al., 2015).

Environmental challenges such as diet, physical exercise, toxins, and psychological stress can alter DNA methylation in gametes (Donkin and Barrès, 2018). Our group and others have shown that the DNA methylation signature of human spermatozoa is amenable to lifestyle influence, such as weight loss and endurance training (Denham et al., 2015; Donkin and Barrès, 2018; Donkin et al., 2016; Ingerslev et al., 2018). Environmentally induced epigenetic change in gametes, therefore, constitutes a plausible mechanism by which environmental factors before conception may influence the phenotype of the next generation. However, the functional consequences of environmentally induced spermatozoal DNA methylation on embryo development have not been elucidated.

As spermatozoa are haploid, methylation levels at a single CpG site directly correspond to the fraction of cells carrying a methyl group on that cytosine. Thus, environmentally induced DNA methylation changes at the single CpG dinucleotide that are smaller than 100% imply that only a fraction of cells in the ejaculate carry that change. For example, a modest increase in methylation on a single CpG site detected in a whole ejaculate from 20% to 40% implicates that an additional 20% of spermatozoa carry a methyl group on that cytosine residue. It is currently unknown if environmentally induced DNA methylation changes across the genome are carried by a sub-population of spermatozoa or are spread across the ejaculate population in a mosaic-like fashion. This information will help to increase our understanding of the mechanisms by which the sperm epigenome is remodeled in response to environmental factors and may have implications for the field of assisted reproduction where selection of single spermatozoa is performed.

Recent developments in single-cell sequencing technologies have allowed for greater exploration of human development (Wen and Tang, 2019). Techniques such as single-cell reduced representation bisulfite sequencing (scRRBS) or post-bisulfite adaptor tagging (PBAT)-based single-cell bisulfite sequencing (scBS-seq) have now been used to profile the DNA methylome of human preimplantation embryos, as well as oocytes and sperm (Guo et al., 2015; Smallwood et al., 2014a; Zhou et al., 2019; Zhu et al., 2018). Although spermatozoal single-cell DNA methylation sequencing has been performed previously in two individuals (Guo et al., 2015; Zhu et al., 2018), it has been done to validate the technical feasibility of the method and to serve as a macroscopic comparison to the oocyte. A detailed DNA methylation profiling of spermatozoa has not been conducted to date.

Here, we performed an adapted PBAT scBS-seq analysis of human sperm cells at single-cell and single-base resolution to map the distribution of DNA methylation marks across cells of a single ejaculate as well as between individuals. We confirm DNA methylation differences between spermatozoa from lean and obese individuals and establish that spermatozoa within one ejaculate have distinct patterns and may harbor marked defects in methylation at specific imprinted loci. Our results provide novel insight into the heterogeneity of sperm DNA methylation in human ejaculates.

## RESULTS

### Post bisulfite adaptor tagging single-cell sequencing of spermatozoa

To compare the epigenetic profile of single spermatozoa from lean and obese individuals, we adapted our PBAT based scBS-seq to circumvent the dense packaging of spermatozoa (Clark et al., 2017; Smallwood et al., 2014b; Smith et al., 2012) by including an additional lysis step and a longer bisulfite conversion (see Methods) (Clark et al., 2017)(Figure 1A). In total, we sequenced 87 spermatozoa from 4 lean and 4 obese individuals at a sequencing depth averaging 15.93 million (M) 100 base-pair (bp) paired-end reads (range, 1.5M to 216.2M reads) (Table S1 and Figure 1B). Although our mapping rate was lower than somatic cells, it was consistent with a previous study using PBAT scBS-seq in spermatozoa (Table S1 and Figure 1C) (Clark et al., 2017; Smallwood et al., 2014b; Zhu et al., 2018). (Figure 1C). Our results are highly correlated with another dataset of spermatozoa (Zhu et al., 2018), but not with DNA methylation data of single oocytes (Zhu et al., 2018) (Figure 1D and Figure 1E). Plotting DNA methylation signals of single cell spermatozoa and oocytes by multi-dimensional scaling (MDS) also showed that this profiling of spermatozoal DNA methylation is specific and consistent with the literature (Figure 1F and Figure S1A). In our dataset, we report global DNA methylation of ∼65.5 %, which is lower than earlier WGBS datasets (∼75.6% in (Guo et al., 2014) and 67.4-72.4% (Molaro et al., 2011)) (Figure 1G). This is suspected to arise from the selection of regions that are present during the PBAT sc-BS-seq protocol, as CpG site comparisons showed similar results. Overall, our analysis suggests that our sequencing data are of a comparative quality to similar published data.

**Figure 1.**
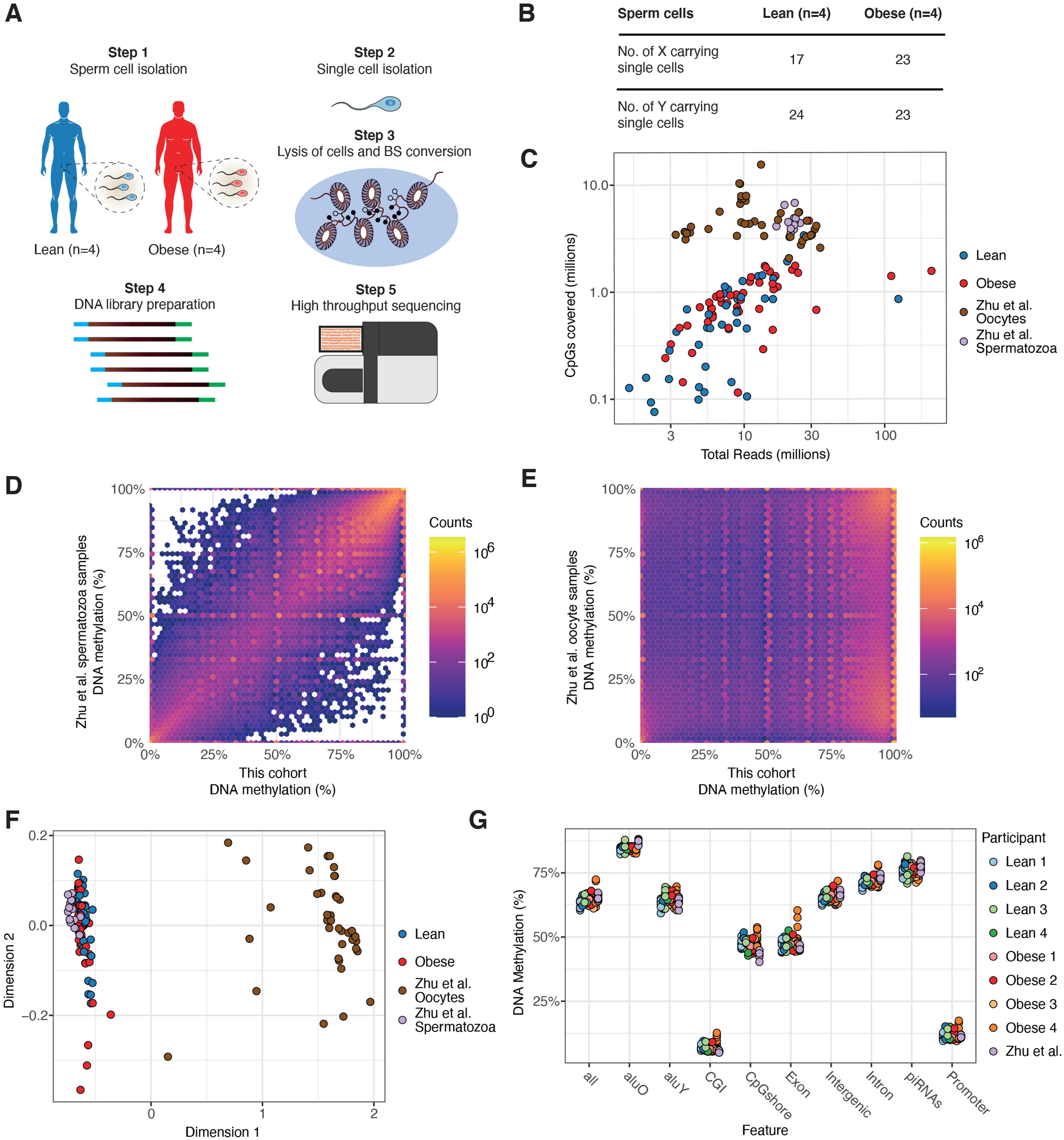
**Benchmark of DNA methylation results from single cell spermatozoa** (A) Schema describing the PBAT scBS-seq method and study participants. Blue and red represents lean and obese respectively. (B) Number of spermatozoa analyzed in each group and based on sex chromosome. All spermatozoa were isolated from the semen sample with a swim-up method to isolate motile spermatozoa and then sorted into a 96-well via flow cytometry using Hoechst 33258 to stain for DNA. (C) Number of CpGs covered by scBS-seq as a function of total mapped sequences compared to a reference dataset (Zhu et al., 2018). (D) DNA methylation levels comparing spermatozoa from our cohort with spermatozoa from Zhu et al. (Zhu et al., 2018). (E) DNA methylation level comparing spermatozoa from our cohort with oocytes from Zhu et al (Zhu et al., 2018). (F) MDS analysis of the DNA methylation profile of single cells compared to spermatozoa and oocytes from Zhu et al (Zhu et al., 2018), with the removal of four outliers described in Zhu et al (Zhu et al., 2018). (G) Average DNA methylation of single spermatozoa based on region and subject.

Because of the high concordance between the results of our study cohort and the results obtained by Zhu et al., we included these data on 23 sperm cells in subsequent analyses to increase power. The single-cell data from Zhu et al. all originated from the same individual and have not been thoroughly used to investigate the role of DNA methylation in single-cell sperm.

### Autosomal differences in DNA methylation in X- and Y-carrying spermatozoa

Given the plethora of sex-specific F1 phenotypes that have been reported in intergenerational inheritance studies (Barbosa et al., 2016; Ng et al., 2010), we sought to determine if X-carrying spermatozoa harbor distinct DNA methylation patterns compared to Y-carrying spermatozoa. “Sex” of the spermatozoa was inferred by mapping reads to X and Y chromosomes (Figure 2A and Table S1). Similar to bulk sequencing, X chromosomes carried more methylation than Y chromosomes (Figure 2B). The autosomal chromosomes of X- and Y-carrying cells showed similar global DNA methylation profiles (Figure 2C and Figure S2A). To further determine the association between the presence of X or Y chromosomes on autosomal chromosome DNA methylation, we plotted the spermatozoa autosomal DNA methylation signature of each sub-population on an MDS plot. We found no clear separation of samples by sex chromosomes (Figure 2D). Analysis of DNA methylation of autosomal chromosomes showed no difference in X- and Y-carrying spermatozoa (Figure 2E). A similar result was observed when plotting DNA methylation of X vs. Y chromosomes on a scatter plot, although this analysis revealed small potential differences (Figure S2A). To further compare DNA methylation in X- and Y-carrying spermatozoa at higher resolution, we performed a comparison at specific genomic regions: 1) gene promoters, 2) Coding Sequences (CDS), and 3) at CpG dense regions, defined as stretches where CpGs are clustered with no more than 100 base pairs apart (Table S2). We found two promoters with higher methylation in the male spermatozoa, one for Tubulin Polyglutamylase Complex Subunit 1 (*TPGS1*) and one for Zic Family Member 2 (*ZIC2*). No differential methylation was detected for CDS or CpG dense regions (Figure S2B and Figure S2C). Overall, our results indicate that DNA methylation of autosomal chromosomes is distinct in X- and Y-carrying spermatozoa, although these differences appear to be modest.

**Figure 2.**
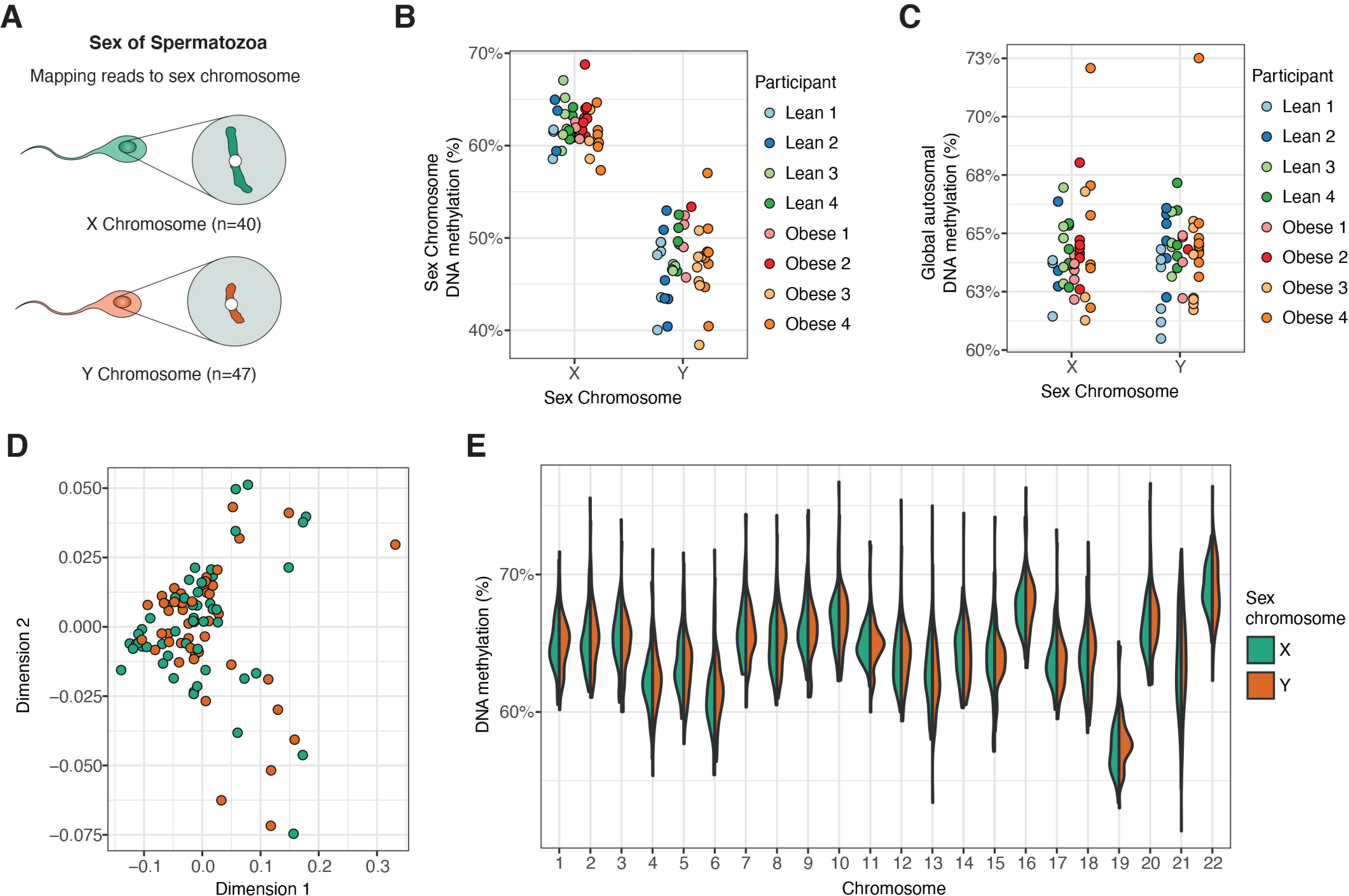
**Distinct DNA methylation landscape in sperm with X and Y chromosome** (A) Schema of X and Y chromosome carrying spermatozoa. Green and orange denotes X chromosome carrying and Y chromosome carrying spermatozoa. (B) Global DNA methylation level of the X and Y chromosome single spermatozoa. (C) Global DNA methylation of autosomal chromosomes in X and Y chromosome single spermatozoa. (D) MDS analysis of the DNA methylation profile of single cells based on sex chromosomes. (E) Bean plot of autosomal DNA methylation based on chromosome and sex chromosome. Kolmogorov–Smirnov test showed no difference in chromosome DNA methylation of X and Y carrying spermatozoa.

### Extensive imprinting defects exist in subpopulations of spermatozoa within one ejaculate

Next, we examined DNA methylation patterns at imprinted genes in single spermatozoa. While we confirmed that, at the single cell level, maternally (*PEG3*) and paternally (*H19*) methylated imprinted genes had the expected DNA methylation patterns as bulk sequencing would predict (Figure S2A and Figure S2B), we detected two types of single cell imprinting defects. The first type is where a single CpG site was abnormally methylated, for example, at the Insulin-Like Growth Factor 2 (*IGF2*) locus (1,391 base stretch) (Figure 3A). The second type is where single cells carry extensive stretches of DNA methylation defects, with neighboring CpGs carrying similar hypo- or hypermethylation, as seen at the imprinted region near the Insulin-Like Growth Factor 2 Receptor (*IGF2R*) locus (1,002 base stretch) (Figure 3B). As we observed specific sperm cells carrying imprinting defects at long extensive stretches, we wondered if these cells might carry imprinting defects in other regions (Figure 3C). We found that extensive imprinting defects were mainly present at only one imprinted region or with only modest imprinting defects (< 50% DNA methylation change of the CpGs in the imprinted region), however, a minority of cells carried extensive defects at several regions. For example, one cell from the “Obese 4” participant had aberrant hypermethylation at the maternally imprinted genes *IGF1R (1,045 base stretch)*, *GPR1-AS (2,477 base stretch),* and *GNAS XL (2,557 base stretch)*, and one cell from the “Lean 3” participant had hypermethylation defects at maternally imprinted genes *CXORF56 (1,095 base stretch)*, *GRB10 (2,585 base stretch),* and *MKRN3 (5,409 base stretch)* (Figure 3C). Thus, our data show that both local and more global alterations of DNA methylation patterns at imprinted regions exist in spermatozoa within the same ejaculate and suggest that these imprinting defects are not caused by a general imprinting failure and may thus constitute large epimutations.

**Figure 3.**
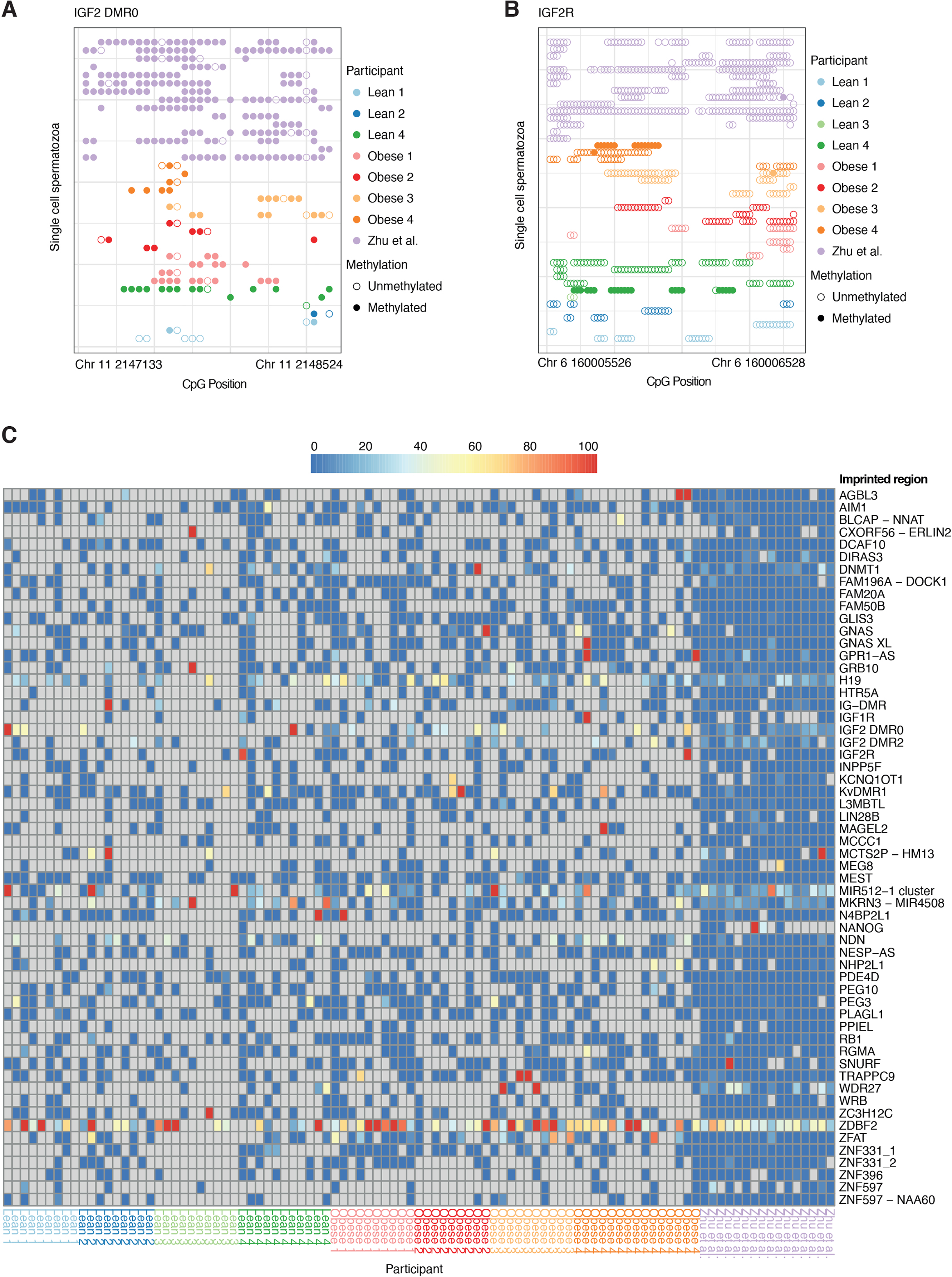
**Single spermatozoa carry imprinting DNA methylation defects** (A) DNA methylation of single spermatozoa at the imprinted region *IGF2*. (B) DNA methylation of single spermatozoa at the imprinted region *IGF2R*. (C) Heat map of the DNA methylation patterns at imprinting regions in single spermatozoa.

### Detection of higher DNA methylation variance at specific transposable elements

To identify genomic regions of DNA methylation heterogeneity in single spermatozoa, we clustered cells based on their methylation at individual CpG sites covered in at least 60 single spermatozoa in the analysis. We identified a total of 292 CpG sites and observed that for most CpG sites, the methylation state was similar across all sperm cells, independently of the donor (Figure S4A). However, we did observe that 183 CpG sites had variations in DNA methylation in at least one of the sperm cells investigated (Figure 4A). Clustering of the cells did not reveal any specific subgroups of spermatozoa. As we observed a range of CpG sites in spermatozoa from the data of Zhu et al. (Zhu et al., 2018) that carried a distinct DNA methylation profile, we sought to investigate whether the difference in DNA methylation observed between spermatozoa of the different cohorts and detected epimutations could be caused by nearby SNPs (Zhu et al., 2018). We found that some of the regions had nearby SNPs, but overall, these did not affect DNA methylation at the detected proximal CpG sites (Figure S4B). Clustering sperm based on methylation levels of CpG dense regions (1,861 clusters covered by a minimum of 60 cells) revealed a similar pattern to that of individual CpG sites, supporting the finding that variations in sperm cells are uniformly distributed (Figure S4C). To determine if specific genomic regions exhibit DNA methylation heterogeneity, we measured the standard deviation of DNA methylation based on the mean methylation level at each CpG region (Figure 4B). Whereas regions at near 0% or 100% methylation displayed consistent low variation, regions with intermediate methylation (25%-75%) contained regions with both low and high variation (Figure 4B). Specifically, for the intermediately methylated regions of high variation, we found that they were more often located far away from the transcription start site, corresponding to being relatively more present in distal intergenic regions and introns, while less present in promoter regions (Figure 4C and Figure S4D). When investigating regions with extensive DNA methylation variation, we found that *young* Alu transposable elements exhibited a particularly high level of DNA methylation variation. Stratifying DNA methylation carried by specific species of transposable elements, we found that single cell DNA methylation variation was higher in *young* Alu elements as well as certain species of LINE-1 (L1) retroelements and “short interspersed nuclear element, variable number of tandem repeats, and Alu composite” (SVA) (Figure 4D, Figure S4E and Figure S4F). As sequencing depth and average DNA methylation level could be responsible for the higher variation found in the younger transposable regions, we investigated the number of sequencing counts for each type of Alu element (Figure S4G). We found that the Alu with the lowest genomic abundance and therefore the lowest number of counts showed the highest variation (Figure S4G). We found that even though Alu, AluJ, and AluS elements had differential variation, their variation did not seem to be affected by their DNA methylation level (Figure S4H). Comparably, Zhu et al. (Zhu et al., 2018) observed higher variation of methylation at young transposable elements. Since young Alu elements are less abundant than old Alu elements, we down-sampled the coverage of the old Alu elements to match that of the young Alu elements (Figure S4I), which indicates that higher variation of *young* Alu transposable elements is biologically true rather than an artifact of sequencing bias.

**Figure 4.**
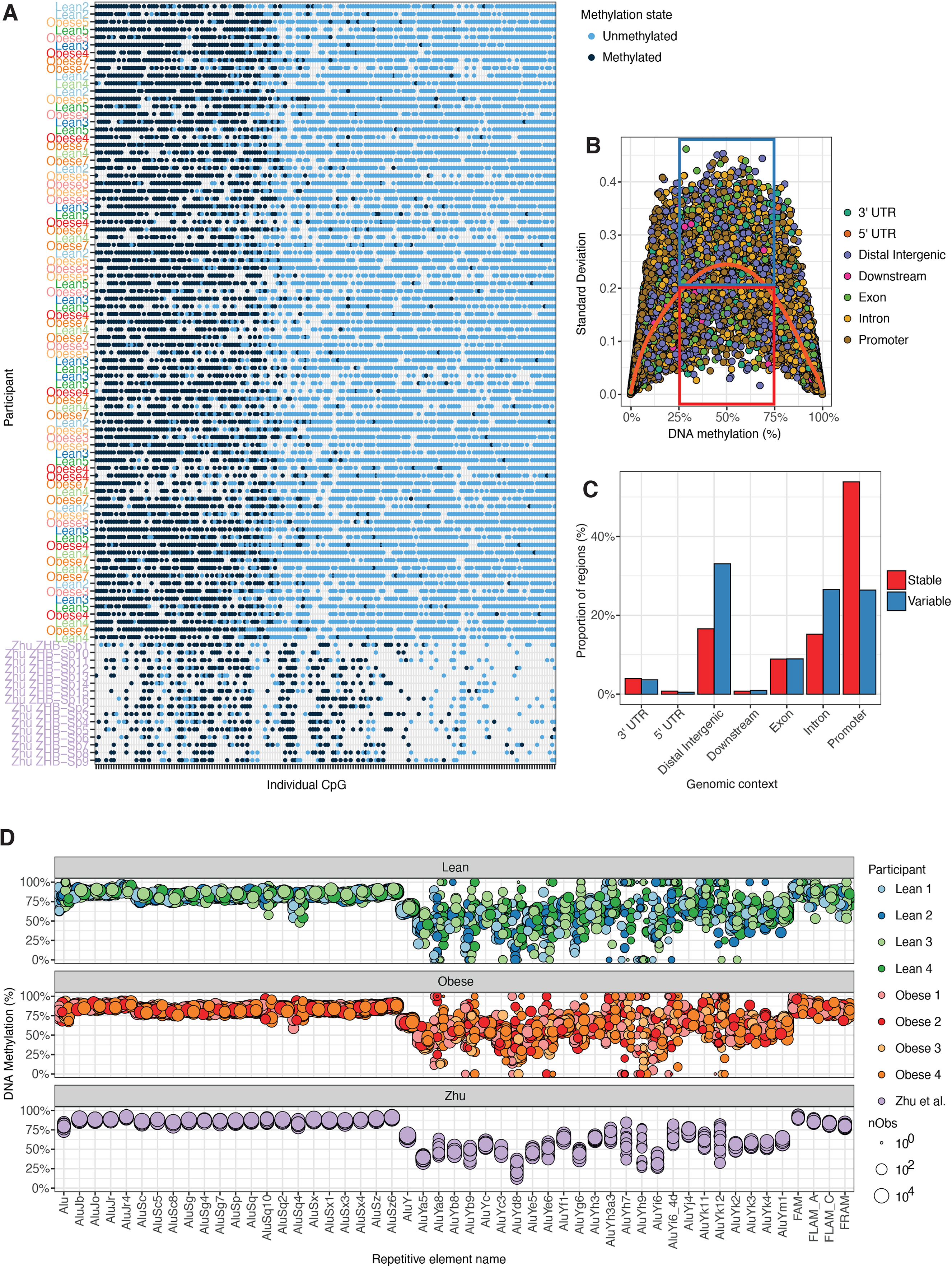
**Heterogeneity of DNA methylation at transposable regions in single spermatozoa** (A) Plotting of the DNA methylation levels at single CpG sites found in at least 60 single spermatozoa and where a variable DNA methylation was found in at least one single spermatozoa. Each line represents a single cell, and each dot represents a given CpG. The location of the CpGs is spread across the genome. (B) DNA methylation variation in single cell DNA methylation clustered by region. (C) Location of stable and variable DNA methylation clusters based on genomic regions. (D) Single cell DNA methylation level at repetitive elements. Each dot represents the DNA methylation level in a specific single cell.

### Single-cell differences in DNA methylation between lean and obese individuals

Previously, we found that spermatozoa from humans with obesity have distinct DNA methylation profiles compared to those of lean men (Donkin et al., 2016). To determine if epigenetic differences are carried by the entire population in a mosaic-like fashion or by a sub-set of spermatozoa, we analyzed single-cell bisulfite data of spermatozoa from lean and obese individuals, with a mean BMI of 23.6 kg/cm^2^ (22.3 kg/cm^2^ −27.2 kg/cm^2^) and 36.8 kg/cm^2^ (32.7 - 40.9 kg/cm^2^), respectively (Table 1). Visualization of single-cell sequencing data comparing spermatozoa of lean and obese men using dimensional scaling revealed a separation of cells from obese individuals at principal component 6 vs. 2, suggesting modest changes (Figure 5A and Figure S5A). Due to the lack of overlap between individual CpG’s across our single-cell data, we opted to investigate differences in DNA methylation at the feature level, specifically at gene promoters, CDS and CpG dense regions (Table S3). We identified two CDS and two CpG dense regions with significantly different DNA methylation levels between cells from lean and obese men; one CDS region on chromosome 7 near *ASZ1* and one on chromosome 14 near *ENTPD5*, as well as one CpG dense region on chromosome 2, related to *LINC01237*, and one on chromosome 17, located near *PPM1D* (Figure 5C, Figure 5D, Figure S5A and Figure S5B). The cluster analysis for *LINC01237* and *PPM1D* at a single-cell level revealed that even across large regions (2074 and 775 base stretch for *LINC01237* and *PPM1D*, respectively), the differences in DNA methylation that we observed are carried by single cells, in a similar way to what we observed for imprinted genes. (Donkin et al., 2016). Our results suggest that some DNA methylation differences between obese and lean men are carried by a subset of spermatozoa.

**Figure 5.**
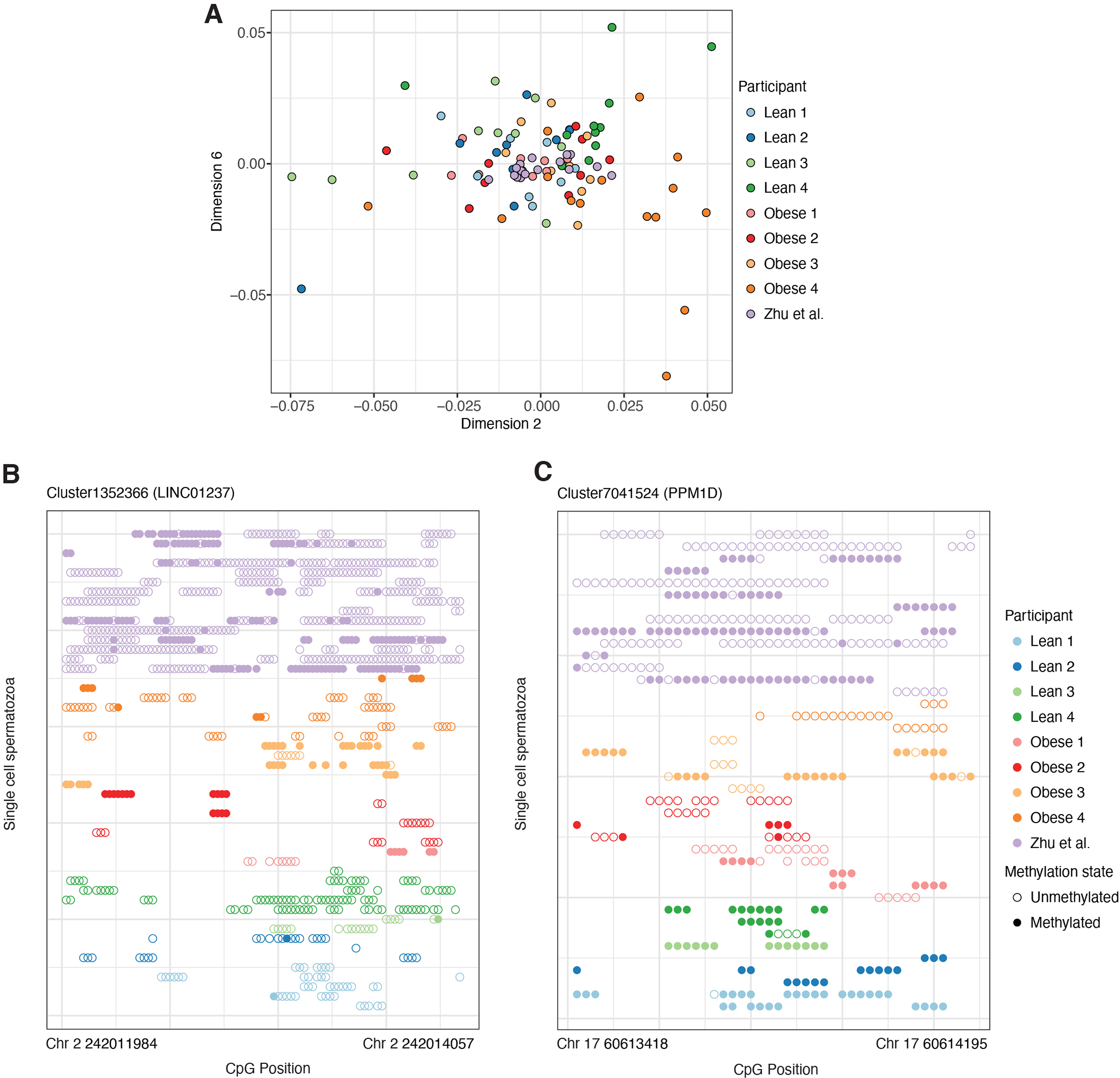
**Differential methylation of single spermatozoa in lean *vs.* obese** (A) MDS analysis of DNA methylation profiles of single cells from lean and obese individuals. (B) DNA methylation of the region *LINC01237* in single spermatozoa from the various participants. (C) DNA methylation of the region *PPM1D* in single spermatozoa from the various participants.

**Table 1.** Clinical characteristics of subjects.

|  | Lean (n = 4) | Obese (n=4) |
| --- | --- | --- |
| Age (yr) | 24.25 [21-26] | 32.21 [23.13-47.17] |
| Weight (kg) | 82.25 [69-98] | 120.2 [97.4-142.7]* |
| Height (m) | 1.86 [1.76-1.93] | 1.80 [1.73-1.93] |
| BMI (kg/m<sup>2</sup>) | 23.62 [22.26-27.15] | 36.79 [32.66-40.91]* |
| Volume (ml) | 3.16 [0.95-5.3] | 2.33 [1.45-2.7] |
| Sperm concentration (million/ml) | 88.50 [20.40-115.60] | 53.06 [6.10-122.70] |
| Total sperm count (million) | 328.68 [19.38-604.20] | 140.04 [8.85-331.29] |
| Motile (%) | 36.82 [8.6-62.42] | 26.87 [17.9-41.01] |
| Total motile sperm count (million) | 136.16 [5.28-377.14] | 37.66 [2.43 - 69.90] |

Data are represented as mean (minimum-maximum).
\*Difference versus lean analyzed with student's t-test.

## DISCUSSION

Here we analyzed single-cell genome-wide DNA methylation data from spermatozoa of single ejaculates from several lean and obese individuals. We identify that within the same ejaculate, spermatozoa carrying X chromosomes have specific DNA methylation profiles compared to those carrying Y chromosomes. We confirm that spermatozoa from lean and obese men display distinct DNA methylation signatures, and we identify that a subset of spermatozoa within one ejaculate carry DNA methylation differences across large regions. Additionally, a subset of spermatozoa from single ejaculates exhibits large imprinting defects independently of BMI. Our results showing distinct epigenetic sub-populations of spermatozoa within an ejaculate have important implications for reproductive biology and assisted reproduction techniques.

Our study demonstrates that PBAT scBS-seq can be used to sequence single-cell spermatozoa and that the DNA methylation patterns obtained with this method are in good agreement across laboratories. However, this sequencing requires very high depth due to the low mapping rate and can only provide insight on a small fraction of the genome, with about 1-5 % of the total amount of CpGs covered in each cell. The fact that we observed a lower global methylation level compared to bulk sequencing suggests that the methylated and more tightly packaged heterochromatin was sequenced at a lower rate (Molaro et al., 2011). One way of solving this issue in future studies could be to use statistical models to predict neighboring CpG methylation. This method has been successfully performed in other cell types where larger datasets are available (Souza et al., 2020). Alternatively, future developments will undoubtedly allow for increased mapping rates and information to be recovered from more CpG sites. For example, bisulfite free methylation methods may prove useful and due to less harsh treatment conditions of DNA, may both increase mapping rate and reduce bias in single-cell sequencing. However, these methods are still undergoing validation in bulk sequencing (Liu et al., 2019). Overall, our results suggest that scBS may be reliably used to detect DNA methylation of single spermatozoa between individuals and even across laboratories. Nonetheless, the method has disadvantages, especially its cost efficiency, which is very low due to the low mapping rate compared to WGBS methods.

As males and females differ only at the genetic level by the sex chromosomes most sexual dimorphisms are assumed to be driven by gene expression differences on autosomal genes (Lopes-Ramos et al., 2020; Oliva et al., 2020; Wijchers and Festenstein, 2011). Single-cell DNA methylation sequencing of spermatozoa allowed, for the first time, the ability to compare if X and Y chromosome DNA methylation differences exist on autosomal chromosomes before conception. We show evidence of altered DNA methylation at two promoter regions. Given that we only analyzed a fraction of the genome, more DNA methylation differences between X and Y chromosome-carrying spermatozoa are likely when studying the whole genome. Interestingly, one of the regions that we identified as differentially methylated is near the gene, *TPGS1*, which has a sex-specific expression pattern in the neonatal mouse cortex/hippocampus before sexual differentiation (Armoskus et al., 2014). Sex-specific DNA methylation differences have been reported in both cord blood and placentas of newborns, as well as in the blood of adults, where it was also linked to an altered gene expression pattern (Gong et al., 2018; Lopes-Ramos et al., 2020; Maschietto et al., 2017; Oliva et al., 2020; Singmann et al., 2015). Understanding autosomal differences will be important for examining the underlying cause of the plethora of sex-specific F1 phenotypes that have been reported in intergenerational inheritance studies (Barbosa et al., 2016; Ng et al., 2010).

There is a growing number of known imprinting disorders, which affect early embryonic development, brain development, male infertility (Hajj et al., 2011; Hattori et al., 2019; Santi et al., 2017) and metabolic adaption later in life (Constância et al., 2004; Tucci et al., 2019). Strikingly, we found that single spermatozoa within an ejaculate have extensive DNA methylation defects at several imprinted regions. This finding is interesting as it has been reported that some patients with imprinting disorders such as Beckwith-Wiedemann syndrome may have more generalized imprinting defects (Bliek et al., 2009; Eggermann et al., 2008). Moreover, imprinted regions are candidates for carrying epigenetic inheritance effects, as these regions escape epigenetic reprogramming after fertilization (Smallwood et al., 2011; Tang et al., 2015; Zhu et al., 2018). Thus, our data demonstrating that some spermatozoa have specific imprinting defects supports a mechanism by which environmentally induced erroneous reprogramming at imprinted regions may not be erased after fertilization. This mechanism may contribute to epigenetic effects in the next generation. It is essential to highlight that the ejaculates that we used in our study were subjected to a swim-up procedure. Therefore, all the spermatozoa that we studied, even those with imprinting defects, were motile and carried the potential to reach the egg. Given the possible dramatic influence of imprinting defects on the development of the embryo and the health of the offspring, our results open a new perspective when considering the selection of spermatozoa, for example, during intracytoplasmic sperm injection.

Our investigations revealed that DNA methylation is mostly homogeneous in spermatozoa, with most DNA methylation variability being equally distributed across the spermatozoa population in a mosaic-like fashion. However, few spermatozoa showed marked imprinting defects. Here, since we have analyzed motile spermatozoa from our swim-up extraction method, our results do not suggest that imprinting defects are carried solely by immotile spermatozoa. Intra-ejaculate differences in sperm DNA methylation have been reported between high-quality and low-quality spermatozoa purified *via* density gradient (Jenkins et al., 2015). An increased coefficient of DNA methylation variation was found in the low-quality subpopulation, and, interestingly, this variation was rarely observed in the same locus between subjects, supporting the idea that the variability in DNA methylation is spread in a random mosaic-like fashion (Jenkins et al., 2015). Our study did find specific regions that presented with a higher variation in DNA methylation. Especially, our finding that young Alu elements have higher variation is intriguing, as these particular regions are associated with spermatozoa fertility potential and constitute one of the few regions that undergo *de novo* methylation in the early developing zygote (Hajj et al., 2011; Zhu et al., 2018). Transposable elements are key drivers of gene expression during early embryonic development, specifically during zygotic gene activation (reviewed in Rodriguez-Terrones & Torres-Padilla 2018)(Rodriguez-Terrones and Torres-Padilla, 2018). In particular, young transposable elements may play an important role in early paternal gene expression (Zhu et al., 2018). While we could not rule out that the detection of increased variance at young Alu elements was not caused by a sequencing bias, it is tempting to speculate that increased variance of young, compared to ancient, Alu elements is due to less efficient DNA methylation maintenance at these regions, thereby leading to phenotypic stochasticity in the next generation.

Obesity is associated with specific DNA methylation patterns in human spermatozoa when sequenced in bulk. Here, we confirmed that at the single-cell level also, obesity is associated with differential DNA methylation in sperm. However, the differentially methylated regions we identified did not overlap with regions previously identified by our group (Donkin et al., 2016). This is likely to be due to the difference in the areas covered as well as the method that we used in the two analyses (Reduced Representation Bisulfite Sequencing of bulk (Donkin et al., 2016) and PBAT scBS-seq (Clark et al., 2017)). Our data indicate that differences between groups may be driven by large hypo- or hypermethylated regions in individual cells, similar to what was observed in imprinted regions. Such single cell specific DNA methylation patterns at specific loci may be due to the processive binding of enzymes establishing and removing DNA methylation (Rulands et al., 2018). It is possible that few cells carry most or all the aberrant methylation patterns, however, the limited coverage inherent to single-cell sequencing did not allow us to determine if the same cells carry aberrant methylation profiles at more than one locus.

Our results allow us to model the mechanisms of epigenetic inheritance, which may involve sperm DNA methylation. In intergenerational epigenetic studies, most of the reported changes in sperm DNA methylation are within the 5-15 % difference range at specific sites (Donkin and Barrès, 2018). In haploid cells like spermatozoa, such figures imply that 5 to 15 spermatozoa out of 100 would have an altered DNA methylation pattern for this specific CpG. Yet, offspring phenotypes described in paternal effects show high penetrance, suggesting that DNA methylation changes driving the phenotypic response may occur at distinct CpG sites, which would independently affect the offspring towards the same phenotype. Given the extensive mosaicism in DNA methylation patterns that we found in spermatozoa within an ejaculate, this mechanism would require that different patterns of CpG methylation variation are integrated into one single phenotype. Such model, by which methylation variation at different genes leads to an integrated phenotypic alteration is plausible in the case of metabolic dysfunction, a phenotype very often measured in epigenetic inheritance studies, as metabolic disorders like obesity and type 2 diabetes are polygenic (Flannick and Florez, 2016; Kong et al., 2012; Manolio et al., 2009; Morris et al., 2012).

From animal studies, it is clear that DNA methylation is essential for spermatogenesis and sperm maturation, as shown by knock-out models of DNA methyltransferases and treatment of animals with the hypomethylation agent 5-azacytidine (Oakes et al., 2007; Takashima et al., 2009; Vasiliauskaitė et al., 2018). Besides a decreased ability to develop mature spermatozoa, one study showed a reduced capacity of the developed spermatozoa, with altered DNA methylation, to reach the blastocyst stage of embryo development (Vasiliauskaitė et al., 2018). In humans, DNA methylation predicts embryo quality (Aston et al., 2015; Jenkins et al., 2016). Correspondingly a link between DNA methylation patterns in sperm and reproductive capacity has been established in obese men, where survival at the cleavage stage (2-cell stage) was equal to fertile controls, however, after genome activation (blastocyst stage), survival was significantly decreased in embryos fertilized by sperm from obese men (Bakos et al., 2011). In light of the elevated risk of developing imprinting disorders in offspring born from assisted reproduction (Hattori et al., 2019; Henningsen et al., 2020; Johnson et al., 2018; Uk et al., 2018), our results suggest that the subpopulation of spermatozoa carrying imprinting defects may be positively selected by assisted reproduction techniques. Our results provide a substantial addition to our understanding of the mechanisms driving epigenetic inheritance and potentially pave the way for the development of novel assisted reproduction strategies aiming to ensure health in the next generation.

## METHODS

### Sample collection

Single ejaculate semen samples were collected from five lean (BMI 20-25) and five obese (BMI 30-45) male subjects in the age range of 20-36 years. Exclusion criteria were daily intake of prescription medicine, smoking, and testicular disease. All participants were controlled for testicular abnormalities by anamnesis. Sperm ejaculates were delivered after 3-5 days of abstinence and stored at 37°C for 30 minutes for liquefaction of the samples. Basic semen parameters, including volume, concentration, and motility were inspected by manual counting on a phase contrast microscope according to WHO guidelines (WHO, 2010) at room temperature (22°C). The human study was approved by the regional ethical committee for the Capital Region of Denmark (Videnskabsetisk Komité, Kongens Vænge 2, 3400 Hillerød, Denmark, Journal no. H-17041722). All participants provided written informed consent before participating in the study.

### Isolation of motile spermatozoa

A swim-up procedure was performed to isolate motile spermatozoa from non-motile spermatozoa and somatic cells. Before this, volume of ejaculates was determined by wide-bore volumetric pipetting. As previously described (Donkin et al., 2016), 0.5 ml of semen was overlaid with 2 ml of swim-up media, comprising of Earle’s Balanced Salt Solution (Sigma-Aldrich, Germany) with 3.2 mg/ml Human Serum Albumin (Sigma-Aldrich, Germany) and 25 mM Hepes (Sigma-Aldrich, Germany) in round-bottom tubes and incubated for 2 hours at 37°C with 5% CO2 at a 45° angle (Donkin et al., 2016). The upper fractions were pooled per ejaculate, centrifuged, resuspended in Phosphate-Buffered Saline (ThermoFisher) and the spermatozoa counted while presence of somatic cells was inspected and noted.

### Single cell sorting

Single spermatozoa were collected by flow cytometry directly into a well of a 96 well plate, containing 5 µl RLT plus (Qiagen) with 1% v/v β-mercaptoethanol (Biorad), using a BD FACSAria™ III. Before sorting, Hoechst 33258 (Sigma) was used to label the DNA content of the cells as previously described (Smallwood et al., 2014a). Briefly, Hoechst were added to the resuspended motile sperm cells, in a concentration of 5 µg/ml for 10 minutes. After staining, the sperm were washed 2 times before flow cytometry. The sperm cells were stored at −80°C until required for library preparation. Negative controls were lysis buffer alone and were prepared and processed in parallel with single-cell samples.

### Single-cell library preparation

Collected cells were thawed and the following solution was added: 1 µl 10% SDS, 1 µl 0.1M DTT, 2 µl Proteinase K (800 units/µl, Sigma), 0.01µl 1M CaCl_2_ and 0.99µl H_2_0. The cells were then incubated for 6 hours at 50 °C for complete lysis. Bisulfite conversion was performed on cell lysates using the EZ DNA Methylation-Direct kit (Zymo). Compared to our original protocol, we modified the conversion to include an extra 98 °C heating for 3 min, 38 min into the conversion step (Clark et al., 2017): i.e. incubation was 98C 8min, 64C 30min, 98C 3min, 64C 3 hours. Purification was performed as previously described (Clark et al., 2017), and DNA was eluted directly into 39ul of first strand synthesis master mix: 0.4 mM dNTPs, 0.4 µM oligo 1 (a truncated Illumina read 1 sequence followed by six random bases) and 1× Blue Buffer (Enzymatics) before incubation at 65 °C for 3 min followed by a 4 °C pause. 50 U of Klenow exo–(Enzymatics) was added and the samples were incubated at 4 °C for 5 min, +1 °C/15 s to 37 °C, 37 °C for 30 min. Samples were incubated at 95 °C for 1 min and transferred immediately to ice before addition of fresh oligo 1 (10 pmol), Klenow exo–(25 U), and dNTPs (1 nmol) in 2.5 µl of 1x blue buffer. The samples were incubated at 4 °C for 5 min, +1 °C/15 s to 37 °C, 37 °C for 30 min. This random priming and extension was repeated a further three times (five rounds in total). Samples were then incubated with 40 U exonuclease I (NEB) for 1 h at 37 °C before DNA was purified using 0.8× Agencourt Ampure XP beads (Beckman Coulter) according to the manufacturer’s guidelines. Samples were eluted in 49 µl of 0.4 mM dNTPs, 0.4 µM oligo 2 and 1× Blue Buffer. Samples were incubated at 95 °C for 45 s and transferred immediately to ice before addition of 50 U Klenow exo–(Enzymatics) and incubation at 4 °C for 5 min, +1 °C/15 s to 37 °C, 37 °C for 90 min. Samples were purified with the addition of 50ul of water and 80ul of binding buffer (Agencourt Ampure XP beads with the beads removed). After two ethanol washes, beads were resuspended in 50 µl of 0.4 mM dNTPs, 0.4 µM PE1.0 forward primer, 0.4 µM indexed iPCRTag reverse primer, 1 U KAPA HiFi HotStart DNA Polymerase (KAPA Biosystems) in 1× HiFi Fidelity Buffer. Libraries were then amplified by PCR as follows: 95 °C 2 min, 14 repeats of (94 °C 80 s, 65 °C 30 s, 72 °C 30 s), 72 °C 3 min and 4 °C hold. Amplified libraries were purified using 0.8× Agencourt Ampure XP beads, according to the manufacturer’s guidelines, and were assessed for quality and quantity using High-Sensitivity DNA chips on the Agilent Bioanalyzer, and the KAPA Library Quantification Kit for Illumina (KAPA Biosystems). Pools of 12–14 single cell libraries were prepared for 100-bp paired-end sequencing on a HiSeq2500 in rapid-run mode (2 lanes/run).

### Bioinformatics

Reads were pre-processed by Trim Galore v. 0.5.0_dev. Alignment, deduplication, and summarization was performed by Bismark v. 0.20.0 with the --non_directional flag set. Forward and reverse reads were processed separately. Further processing was performed using R and figures were generated using ggplot2. Coverage files generated by Bismark was imported into R. Chromosome M was excluded and CpGs were filtered to only include CpGs covered by a single fragment, not hemi-methylated and a CpG in hg38 reference genome. Forward and reverse reads were aggregated to cell level methylation. In cases where a CpG was covered by both the forward and reverse read, the forward read methylation status was used. CpGs overlapping NCBI dbSNP Build 151 (Sherry et al., 2001) were excluded. Each CpG overlapping a promoter or a CDS was annotated with the gene ID and each CpG was overlapped with a CpG cluster, where a cluster is defined as a stretch of DNA with no more than 100 bp between CpG’s. Cells were annotated as X carrying if there was more than 10 times more reads on the X chromosome than on the Y, and Y carrying the opposite. Data from Zhu et al. (Zhu et al., 2018) was processed in a similar manner, except the processing started at Bismark coverage files. Prior to testing for differential methylation, all cells from the same individual were summed up to create a pseudo-bulk dataset. Differential methylation was tested using edgeR. Differential methylation was tested on CpGs aggregated onto clusters, promoters and CDSs. Only regions with eight or more observations were tested. Heatmap figures comparing our data and the data from Zhu et al. (Zhu et al., 2018) as well as the X vs Y carrying spermatozoa was performed based on averages per cluster. Imprinted region was obtained from (Court et al., 2014) and lifted to hg38 using liftOver, additional regions was obtained from (Fend-Guella et al., 2019). Multidimensional scaling (MDS) plots was generated in two steps, first pairwise distances between all samples were calculated as one minus the simple matching coefficient of the methylation status of CpG cites covered in both samples, next principal component analysis was performed on the distance matrix. For the comparisons between autosomal chromosome DNA methylation for X and Y carrying spermatozoa, all covered CpG’s were used and tested using the Kolmogorov–Smirnov test. Genomic annotations of promoters, introns and exons was obtained from GENCODE v. 31 (Frankish et al., 2018), CpG islands and transposable elements was obtained from the UCSC Table Browser (Karolchik et al., 2004) and piRNAs were obtained from piRNAdb (www.pirnadb.org). CpG sites were considered near a mutation if the read they originated from contained a mismatch relative to the hg38 reference genome.

### Data and code availability

Datasets were deposited in the Gene Expression Omnibus (GEO), with the following GEO accession numbers (GSEXXXX). Single-cell analysis pipeline is accessible at: Github

## STAR METHODS

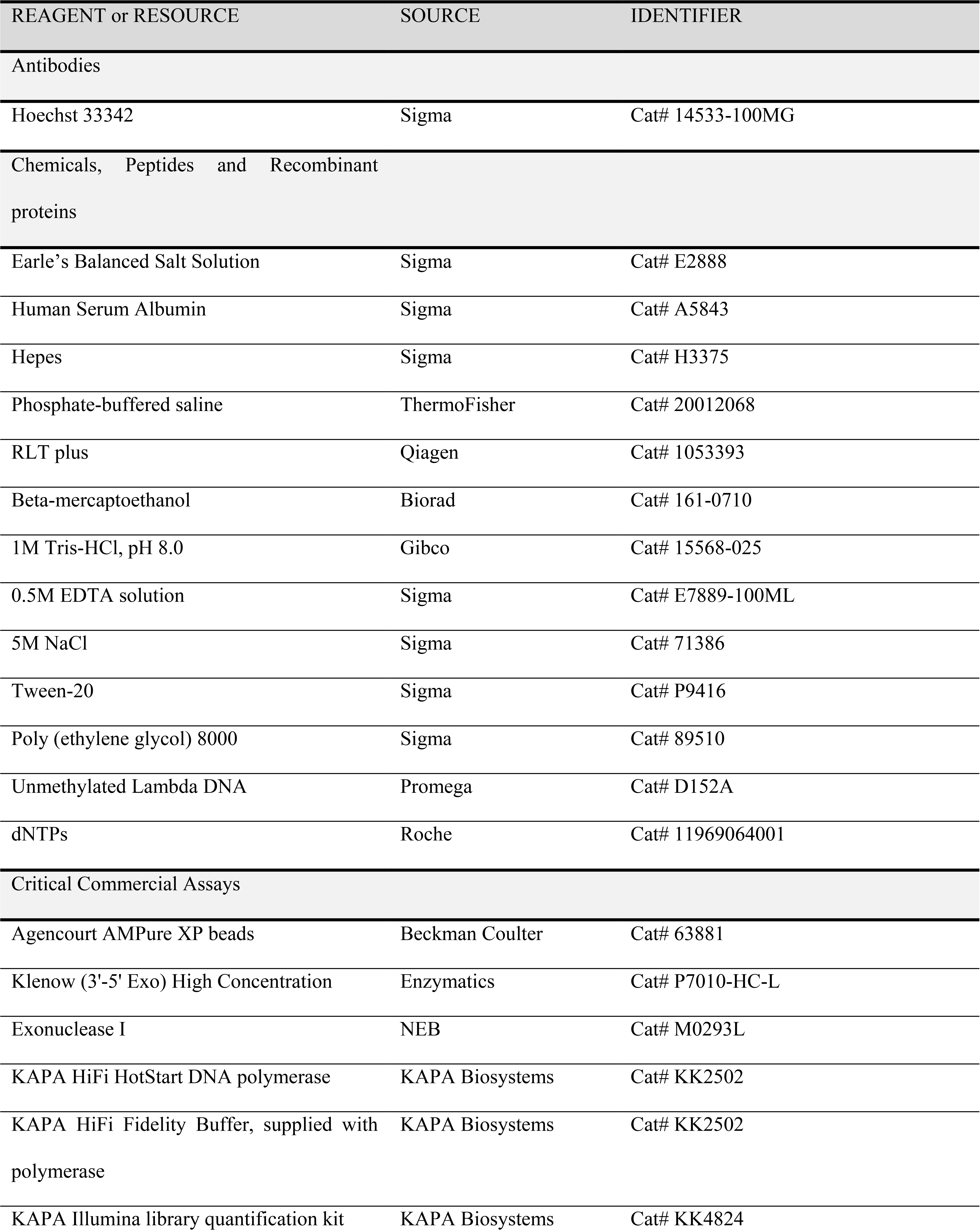
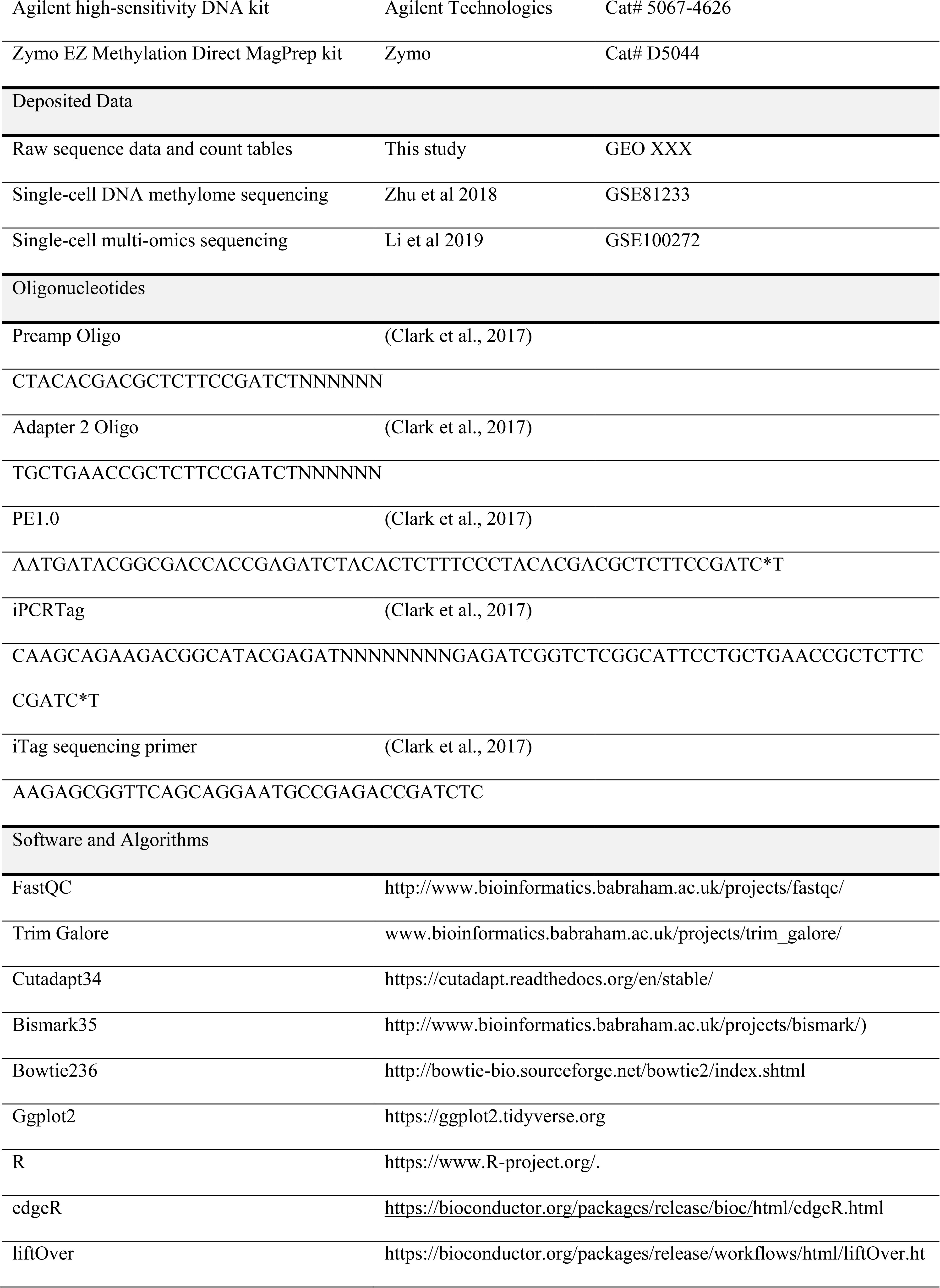
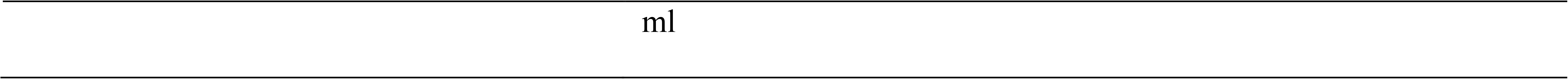

## ACKNOWLEDGMENTS

The authors would like to acknowledge the Flow Cytometry Platform, at the Novo Nordisk Foundation Center for Stem Biology (DanStem) and Center for Protein Research (CPR) for their support and technical expertise. We thank the members of the Barrès Group for their critical reading of the manuscript and helpful comments. The Novo Nordisk Foundation Center for Basic Metabolic Research (http://www.metabol.ku.dk) is an independent research Center at the University of Copenhagen, partially funded by an unrestricted donation from the Novo Nordisk Foundation.

## AUTHOR CONTRIBUTIONS

R.B. and W.R. designed the experiments. E.A. carried out sample collection, swim-up procedure, and flow cytometry isolation. S.C. and E.A. carried out PBAT scBS-seq library preparation. L.R.I performed single cell computational analysis. All authors carried out data analysis and wrote the manuscript.

## SUPPLEMENTARY FIGURE LEGENDS

**Figure S1.**
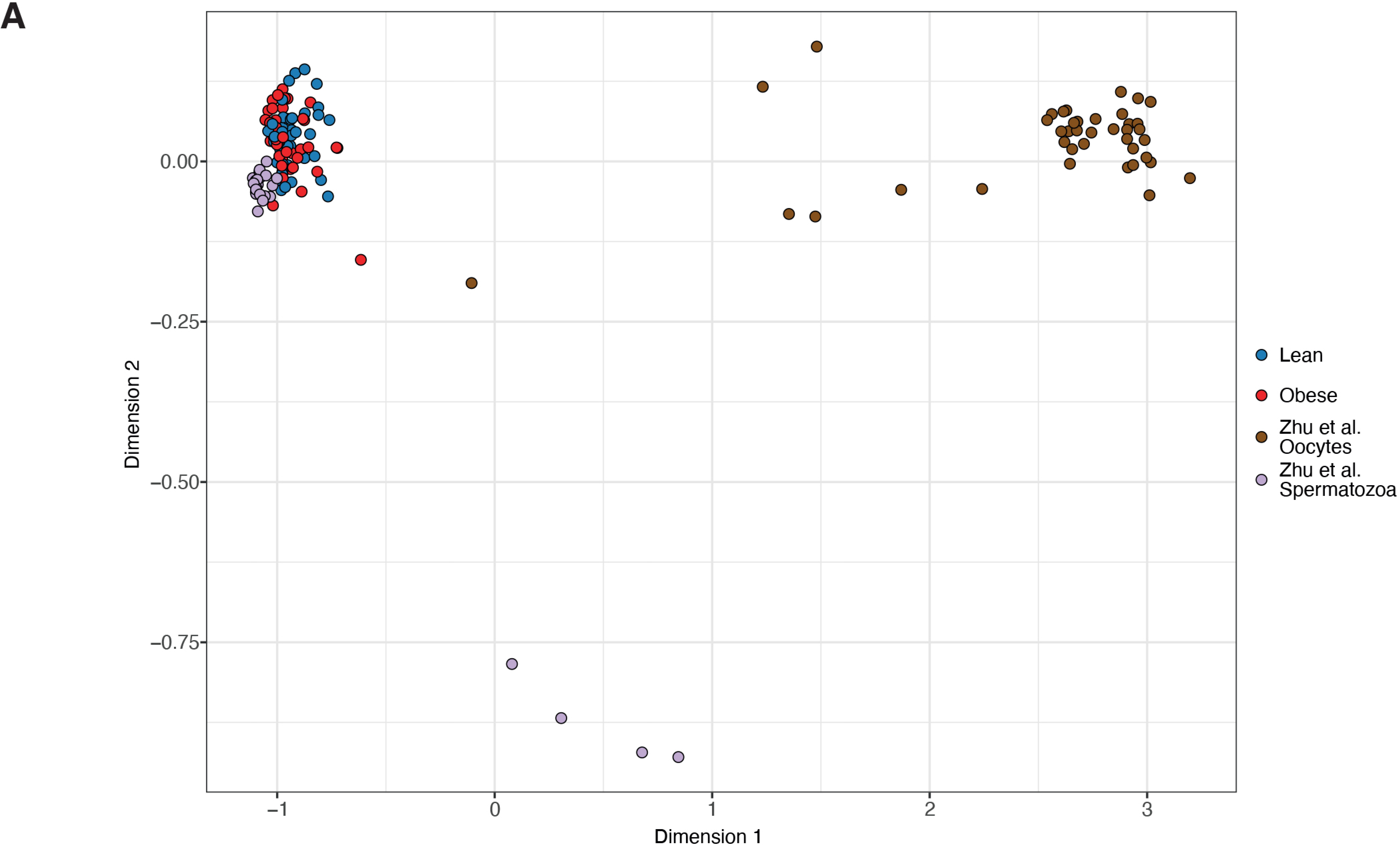
MDS analysis of the DNA methylation profile of single spermatozoa of our cohort compared to spermatozoa and oocytes from Zhu et al.

**Figure S2.**
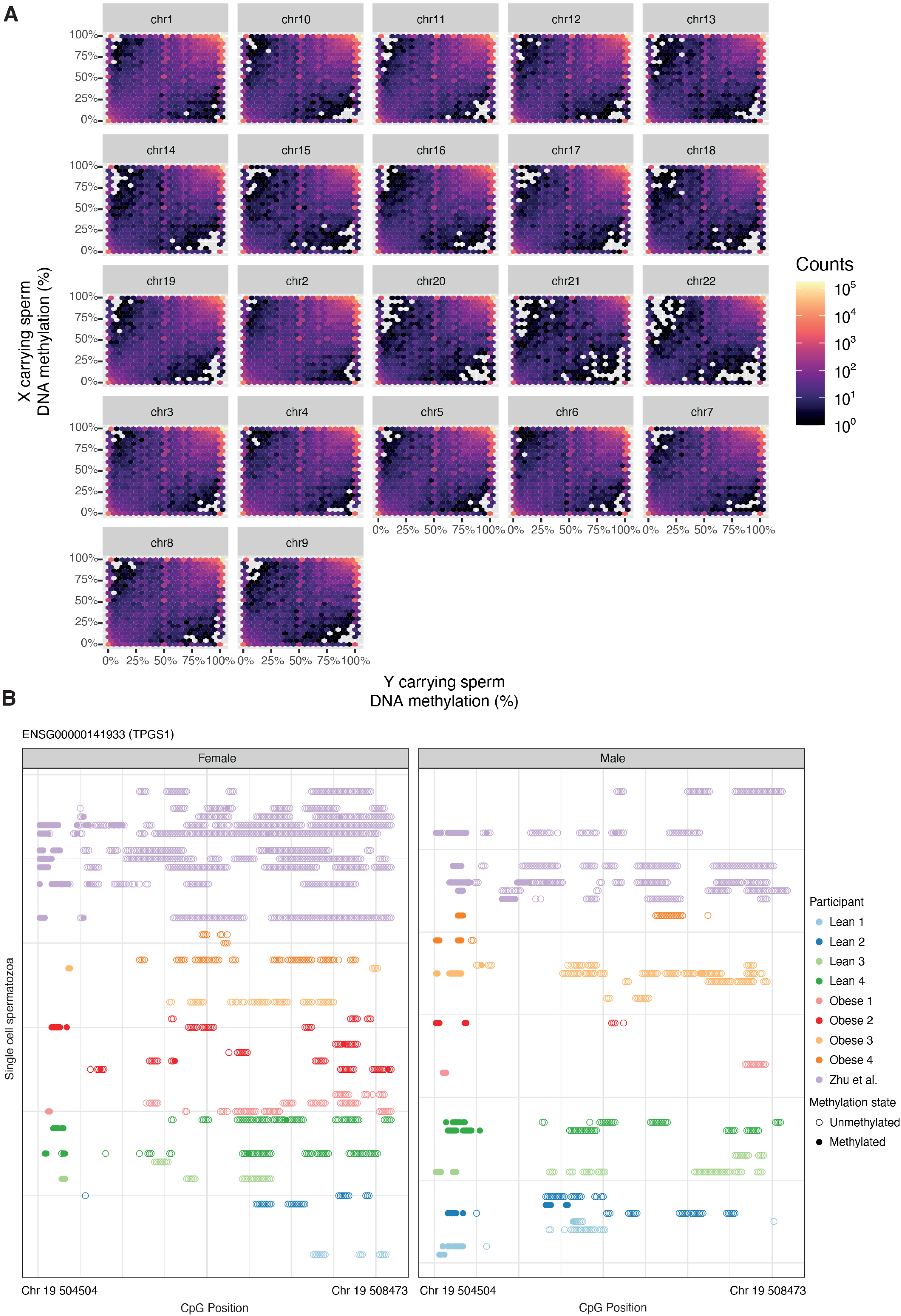
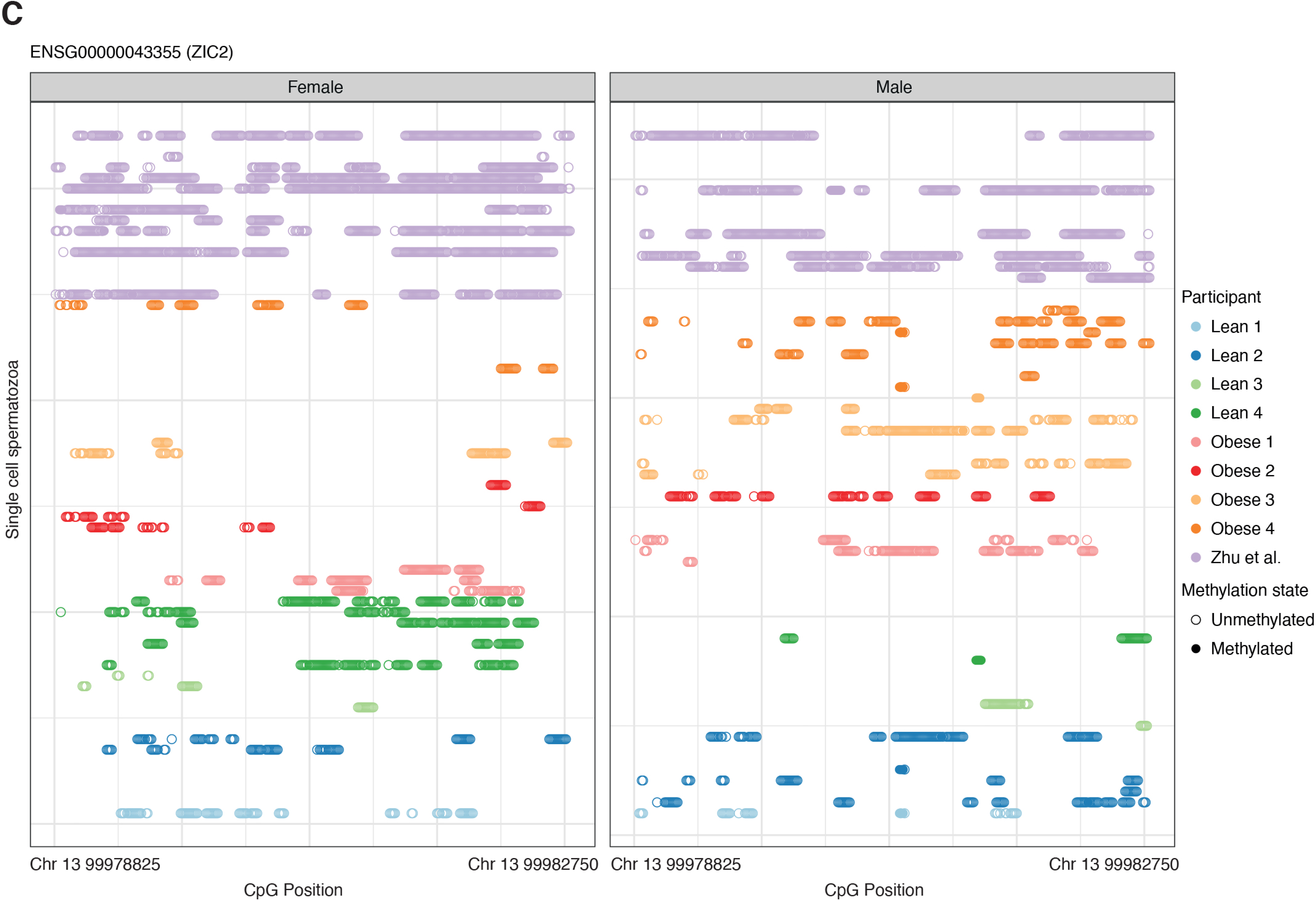
Global and regional DNA methylation differences between X and Y-chromosome carrying spermatozoa. (A) DNA methylation levels of X chromosome compared to Y chromosome. (B) DNA methylation of the differentially methylated *TPGS1* region distributed by X- and Y- carrying chromosomes. (C) Single spermatozoa DNA methylation of the differentially methylated ZIC2 region distributed by X and Y carrying chromosomes in single cells of subjects.

**Figure S3.**
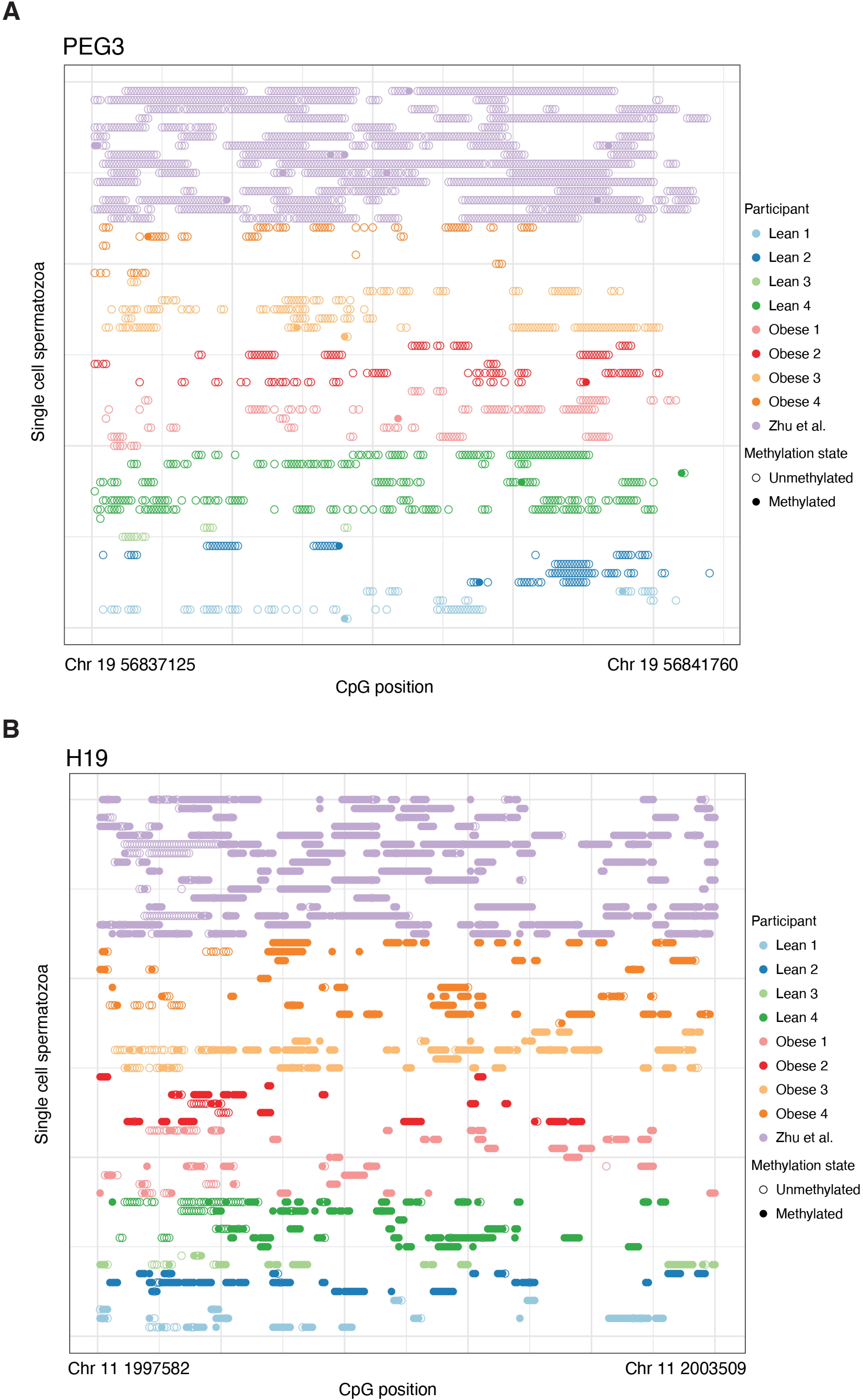
Single spermatozoa DNA methylation at control imprinted regions. (A) Single spermatozoa DNA methylation of the maternally imprinted region PEG3. (B) Single spermatozoa DNA methylation of the paternally imprinted region H19.

**Figure S4.**
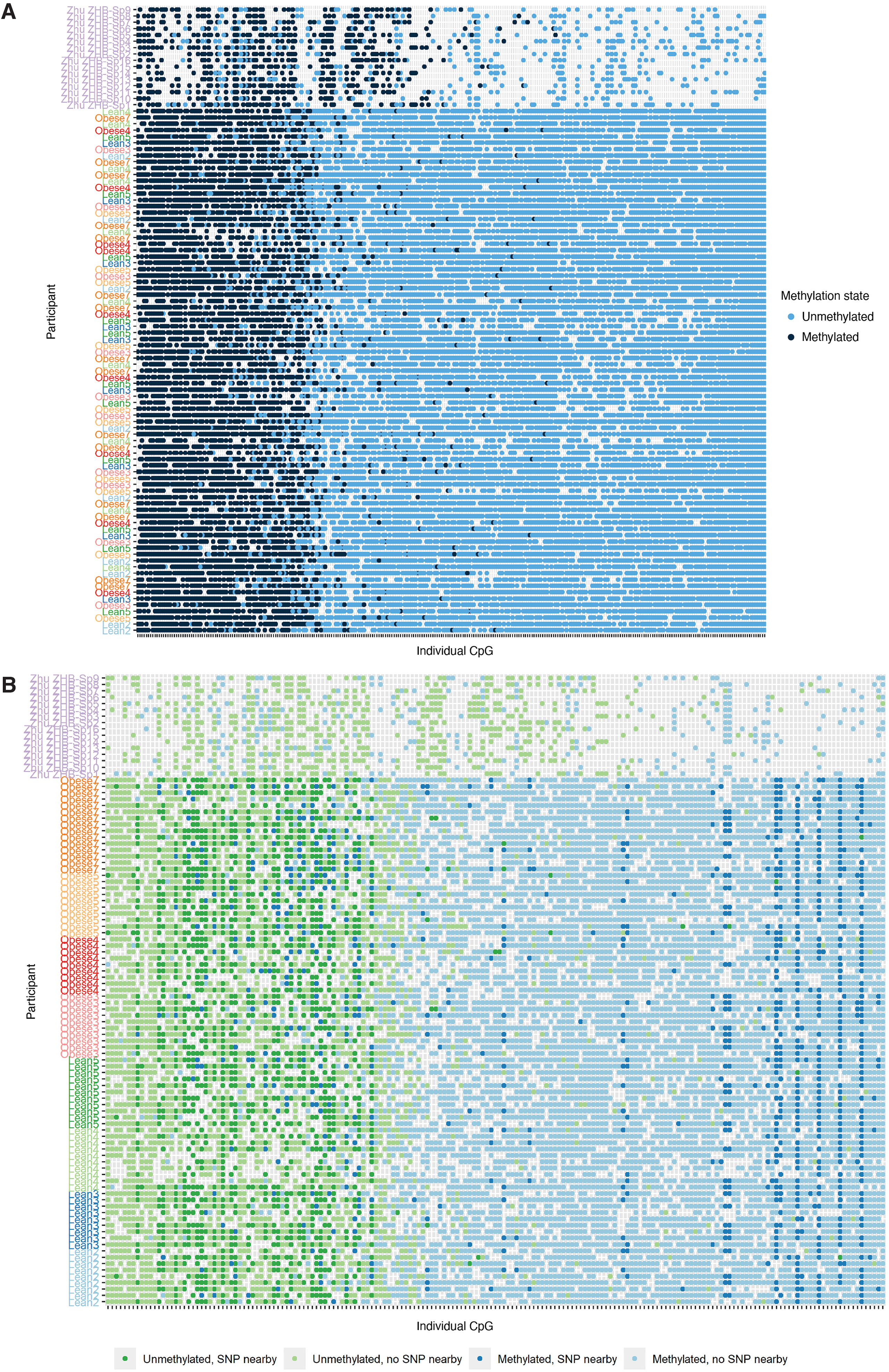
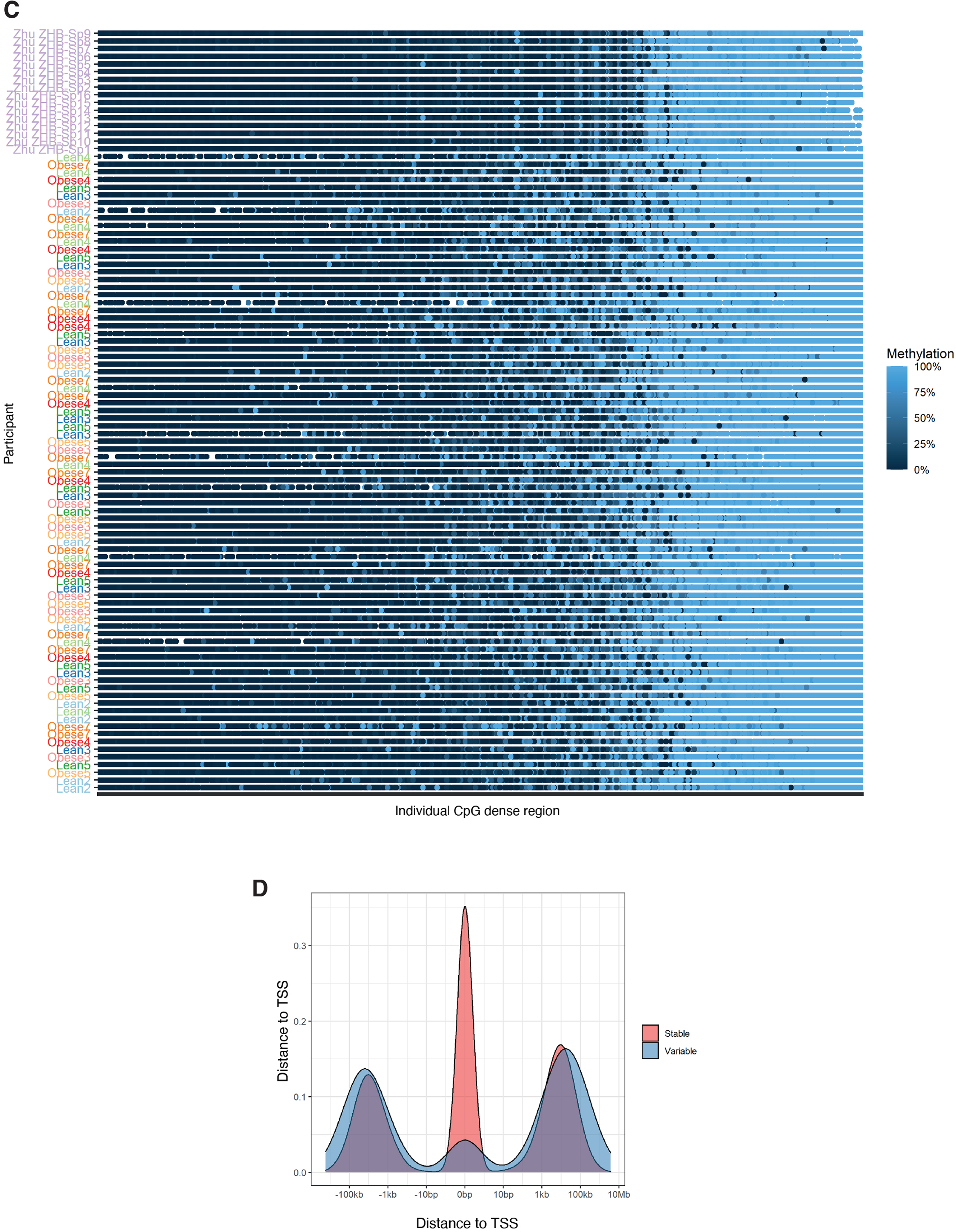
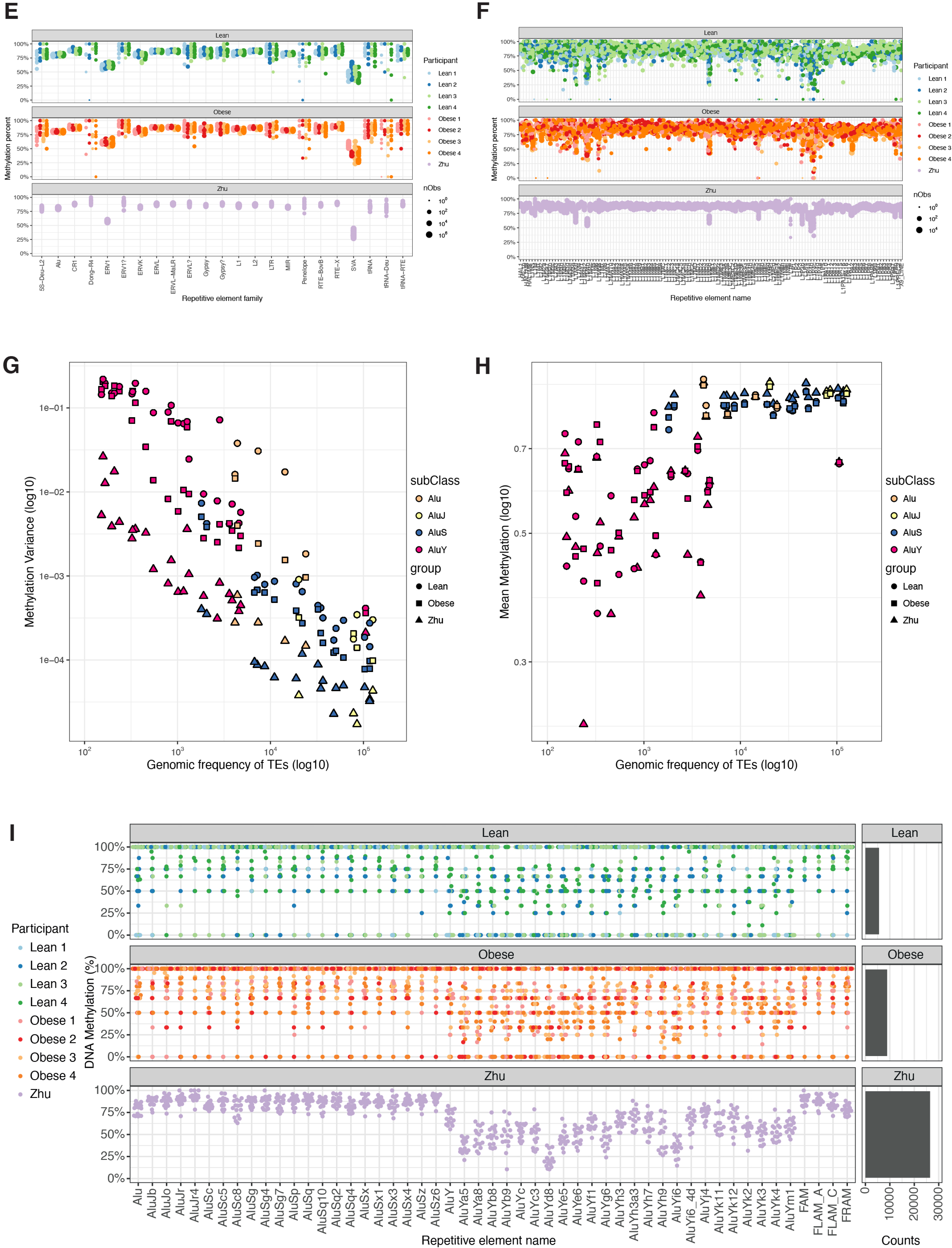
Single-cell DNA methylation heterogeneity of spermatozoa at transposable regions of younger origin. (A) DNA methylation at single CpG sites found in at least 60 single spermatozoa. Each line represents a single spermatozoon, and each dot represents a given CpG. The location of the CpGs is spread across the genome. (B) DNA methylation at single CpG sites found in at least 60 single cells, with a variable DNA methylation in at least one single spermatozoa. Each line represents a single cell, and each dot represents a given CpG. Each dot is colored by DNA methylation state; green (unmethylated) / blue (methylated), and if a SNP was identified within 100 base pairs of the given CpG, light (SNP nearby) / dark (no SNP nearby). The location of the CpGs is spread across the genome. (C) DNA methylation at CpG dense regions identified in all at least 60 single spermatozoa. Each line represents a single cell, and each dot represents a given methylation state of a CpG cluster. The location of the CpG dense regions is spread across the genome. (D) Location of stable and variable DNA methylation clusters based on distance to transcription start site (TSS). (E) Single cell DNA methylation level at repetitive elements. Each dot represents the DNA methylation level in a specific single spermatozoa. (F) Single cell DNA methylation level at repetitive elements. Each dot represents the DNA methylation level in a specific single spermatozoa. (G) DNA methylation variance of Alu subclasses based on their genomic frequency. (H) Mean DNA methylation of Alu subclasses based on their genomic frequency. (I) Single cell DNA methylation level of Alu subclasses where the coverage of each subclass has been down sampled to match that of the lowest Alu subclass per each participant. Each dot represents the DNA methylation level at specific single spermatozoa.

**Figure S5.**
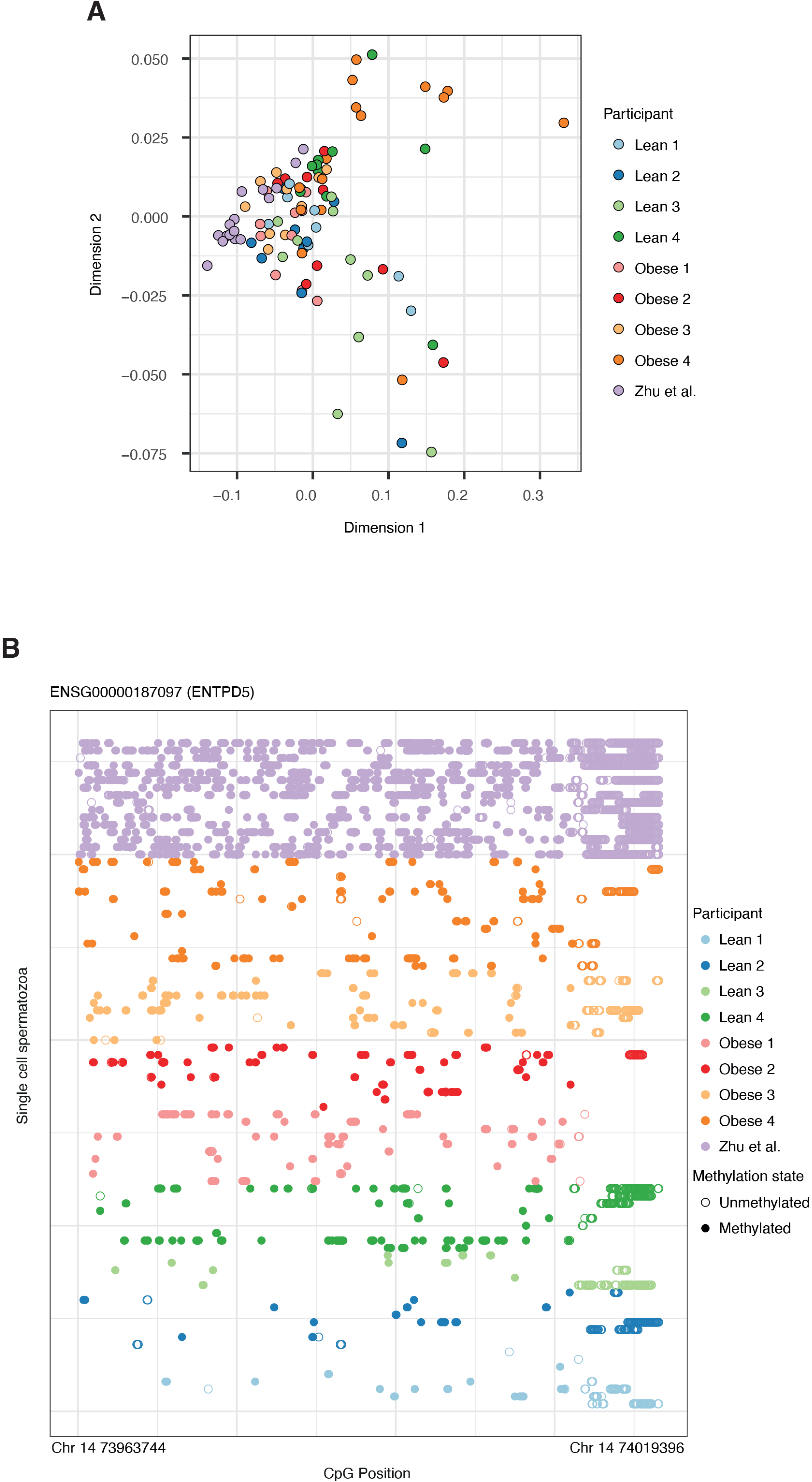
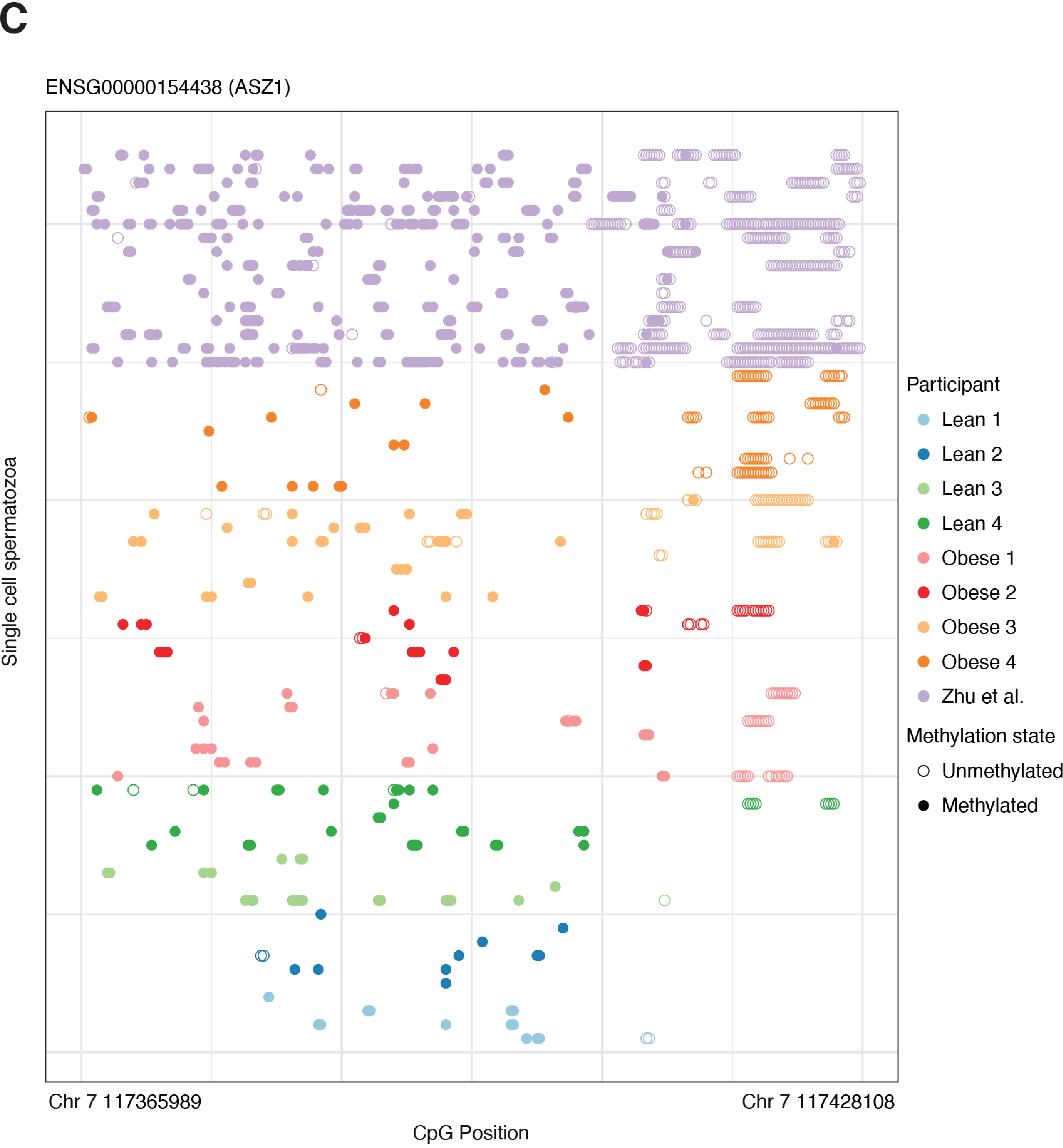
Differentially methylated regions of single-cell spermatozoa from lean and obese individuals. (A) Single spermatozoa DNA methylation of the region *ASZ1* distributed in subjects. (B) Single spermatozoa DNA methylation of the region *ENTPD5* distributed in subjects.

## SUPPLEMENTARY TABLES

**Table S1. Sequencing statistics**

**Table S2. X vs Y chromosome differentially methylated regions**

**Table S3. Differentially methylated regions between lean and obese**

## REFERENCES

Armoskus, C., Moreira, D., Bollinger, K., Jimenez, O., Taniguchi, S., and Tsai, H.-W. (2014). Identification of sexually dimorphic genes in the neonatal mouse cortex and hippocampus. Brain Res 1562, 23–38.

Aston, K.I., Uren, P.J., Jenkins, T.G., Horsager, A., Cairns, B.R., Smith, A.D., and Carrell, D.T. (2015). Aberrant sperm DNA methylation predicts male fertility status and embryo quality. Fertil Steril 104, 1388–1397.e5.

Bakos, H.W., Henshaw, R.C., Mitchell, M., and Lane, M. (2011). Paternal body mass index is associated with decreased blastocyst development and reduced live birth rates following assisted reproductive technology. Fertil Steril 95, 1700–1704.

Barbosa, T. de C., Ingerslev, L.R., Alm, P.S., Versteyhe, S., Massart, J., Rasmussen, M., Donkin, I., Sjögren, R., Mudry, J.M., Vetterli, L., et al. (2016). High-fat diet reprograms the epigenome of rat spermatozoa and transgenerationally affects metabolism of the offspring. Mol Metab 5, 184–197.

Bliek, J., Verde, G., Callaway, J., Maas, S.M., Crescenzo, A.D., Sparago, A., Cerrato, F., Russo, S., Ferraiuolo, S., Rinaldi, M.M., et al. (2009). Hypomethylation at multiple maternally methylated imprinted regions including PLAGL1 and GNAS loci in Beckwith–Wiedemann syndrome. Eur J Hum Genet 17, 611–619.

Clark, S.J., Smallwood, S.A., Lee, H.J., Krueger, F., Reik, W., and Kelsey, G. (2017). Genome-wide base-resolution mapping of DNA methylation in single cells using single-cell bisulfite sequencing (scBS-seq). Nat Protoc 12, 534–547.

Conine, C.C., Sun, F., Song, L., Rivera-Pérez, J.A., and Rando, O.J. (2018). Small RNAs Gained during Epididymal Transit of Sperm Are Essential for Embryonic Development in Mice. Dev Cell 46, 470–480.e3.

Constância, M., Kelsey, G., and Reik, W. (2004). Resourceful imprinting. Nature 432, 53–57.

Court, F., Tayama, C., Romanelli, V., Martin-Trujillo, A., Iglesias-Platas, I., Okamura, K., Sugahara, N., Simón, C., Moore, H., Harness, J.V., et al. (2014). Genome-wide parent-of-origin DNA methylation analysis reveals the intricacies of human imprinting and suggests a germline methylation-independent mechanism of establishment. Genome Res 24, 554–569.

Denham, J., O’Brien, B.J., Harvey, J.T., and Charchar, F.J. (2015). Genome-wide sperm DNA methylation changes after 3 months of exercise training in humans. Epigenomics-Uk 7, 717–731.

Donkin, I., and Barrès, R. (2018). Sperm epigenetics and influence of environmental factors. Mol Metab 14, 1–11.

Donkin, I., Versteyhe, S., Ingerslev, L.R., Qian, K., Mechta, M., Nordkap, L., Mortensen, B., Appel, E.V.R., Jørgensen, N., Kristiansen, V.B., et al. (2016). Obesity and Bariatric Surgery Drive Epigenetic Variation of Spermatozoa in Humans. Cell Metab 23, 369–378.

Eggermann, T., Eggermann, K., and Schönherr, N. (2008). Growth retardation versus overgrowth: Silver-Russell syndrome is genetically opposite to Beckwith-Wiedemann syndrome. Trends Genet 24, 195–204.

Fend-Guella, D.L., Kopylow, K. von, Spiess, A.-N., Schulze, W., Salzbrunn, A., Diederich, S., Hajj, N.E., Haaf, T., Zechner, U., and Linke, M. (2019). The DNA methylation profile of human spermatogonia at single-cell- and single-allele-resolution refutes its role in spermatogonial stem cell function and germ cell differentiation. Mol Hum Reprod 25, 283–294.

Flannick, J., and Florez, J.C. (2016). Type 2 diabetes: genetic data sharing to advance complex disease research. Nat Rev Genet 17, 535–549.

Frankish, A., Diekhans, M., Ferreira, A.-M., Johnson, R., Jungreis, I., Loveland, J., Mudge, J.M., Sisu, C., Wright, J., Armstrong, J., et al. (2018). GENCODE reference annotation for the human and mouse genomes. Nucleic Acids Res 47, gky955-.

Gong, S., Johnson, M.D., Dopierala, J., Gaccioli, F., Sovio, U., Constância, M., Smith, G.C., and Charnock-Jones, D.S. (2018). Genome-wide oxidative bisulfite sequencing identifies sex-specific methylation differences in the human placenta. Epigenetics 13, 01–32.

Guo, H., Zhu, P., Yan, L., Li, R., Hu, B., Lian, Y., Yan, J., Ren, X., Lin, S., Li, J., et al. (2014). The DNA methylation landscape of human early embryos. Nature 511, 606–610.

Guo, H., Zhu, P., Guo, F., Li, X., Wu, X., Fan, X., Wen, L., and Tang, F. (2015). Profiling DNA methylome landscapes of mammalian cells with single-cell reduced-representation bisulfite sequencing. Nat Protoc 10, 645–659.

Hajj, N.E., Zechner, U., Schneider, E., Tresch, A., Gromoll, J., Hahn, T., Schorsch, M., and Haaf, T. (2011). Methylation Status of Imprinted Genes and Repetitive Elements in Sperm DNA from Infertile Males. Sex Dev 5, 60–69.

Hammoud, S.S., Nix, D.A., Zhang, H., Purwar, J., Carrell, D.T., and Cairns, B.R. (2009). Distinctive chromatin in human sperm packages genes for embryo development. Nature 460, 473–478.

Hattori, H., Hiura, H., Kitamura, A., Miyauchi, N., Kobayashi, N., Takahashi, S., Okae, H., Kyono, K., Kagami, M., Ogata, T., et al. (2019). Association of four imprinting disorders and ART. Clin Epigenetics 11, 21.

Henningsen, A.A., Gissler, M., Rasmussen, S., Opdahl, S., Wennerholm, U.B., Spangsmose, A.L., Tiitinen, A., Bergh, C., Romundstad, L.B., Laivuori, H., et al. (2020). Imprinting disorders in children born after ART: a Nordic study from the CoNARTaS group. Hum Reprod 35, 1178–1184.

Ingerslev, L.R., Donkin, I., Fabre, O., Versteyhe, S., Mechta, M., Pattamaprapanont, P., Mortensen, B., Krarup, N.T., and Barrès, R. (2018). Endurance training remodels sperm-borne small RNA expression and methylation at neurological gene hotspots. Clin Epigenetics 10, 12.

Jenkins, T.G., Aston, K.I., Trost, C., Farley, J., Hotaling, J.M., and Carrell, D.T. (2015). Intra-sample heterogeneity of sperm DNA methylation. Mhr Basic Sci Reproductive Medicine 21, 313–319.

Jenkins, T.G., Aston, K.I., Meyer, T.D., Hotaling, J.M., Shamsi, M.B., Johnstone, E.B., Cox, K.J., Stanford, J.B., Porucznik, C.A., and Carrell, D.T. (2016). Decreased fecundity and sperm DNA methylation patterns. Fertil Steril 105, 51–57.e3.

Johnson, J.P., Beischel, L., Schwanke, C., Styren, K., Crunk, A., Schoof, J., and Elias, A.F. (2018). Overrepresentation of pregnancies conceived by artificial reproductive technology in prenatally identified fetuses with Beckwith-Wiedemann syndrome. J Assist Reprod Gen 35, 985–992.

Kaneda, M., Okano, M., Hata, K., Sado, T., Tsujimoto, N., Li, E., and Sasaki, H. (2004). Essential role for de novo DNA methyltransferase Dnmt3a in paternal and maternal imprinting. Nature 429, 900–903.

Karolchik, D., Hinrichs, A.S., Furey, T.S., Roskin, K.M., Sugnet, C.W., Haussler, D., and Kent, W.J. (2004). The UCSC Table Browser data retrieval tool. Nucleic Acids Res 32, D493–D496.

Kong, A., Frigge, M.L., Masson, G., Besenbacher, S., Sulem, P., Magnusson, G., Gudjonsson, S.A., Sigurdsson, A., Jonasdottir, A., Jonasdottir, A., et al. (2012). Rate of de novo mutations and the importance of father’s age to disease risk. Nature 488, 471–475.

Liu, Y., Siejka-Zielińska, P., Velikova, G., Bi, Y., Yuan, F., Tomkova, M., Bai, C., Chen, L., Schuster-Böckler, B., and Song, C.-X. (2019). Bisulfite-free direct detection of 5-methylcytosine and 5-hydroxymethylcytosine at base resolution. Nat Biotechnol 37, 424–429.

Lopes-Ramos, C.M., Chen, C.-Y., Kuijjer, M.L., Paulson, J.N., Sonawane, A.R., Fagny, M., Platig, J., Glass, K., Quackenbush, J., and DeMeo, D.L. (2020). Sex Differences in Gene Expression and Regulatory Networks across 29 Human Tissues. Cell Reports 31, 107795.

Manolio, T.A., Collins, F.S., Cox, N.J., Goldstein, D.B., Hindorff, L.A., Hunter, D.J., McCarthy, M.I., Ramos, E.M., Cardon, L.R., Chakravarti, A., et al. (2009). Finding the missing heritability of complex diseases. Nature 461, 747–753.

Maschietto, M., Bastos, L.C., Tahira, A.C., Bastos, E.P., Euclydes, V.L.V., Brentani, A., Fink, G., Baumont, A. de, Felipe-Silva, A., Francisco, R.P.V., et al. (2017). Sex differences in DNA methylation of the cord blood are related to sex-bias psychiatric diseases. Sci Rep-Uk 7, 44547.

Molaro, A., Hodges, E., Fang, F., Song, Q., McCombie, W.R., Hannon, G.J., and Smith, A.D. (2011). Sperm Methylation Profiles Reveal Features of Epigenetic Inheritance and Evolution in Primates. Cell 146, 1029–1041.

Morris, A.P., Voight, B.F., Teslovich, T.M., Ferreira, T., Segrè, A.V., Steinthorsdottir, V., Strawbridge, R.J., Khan, H., Grallert, H., Mahajan, A., et al. (2012). Large-scale association analysis provides insights into the genetic architecture and pathophysiology of type 2 diabetes. Nat Genet 44, 981–990.

Ng, S.-F., Lin, R.C.Y., Laybutt, D.R., Barres, R., Owens, J.A., and Morris, M.J. (2010). Chronic high-fat diet in fathers programs β-cell dysfunction in female rat offspring. Nature 467, 963–966.

Oakes, C.C., Kelly, T.L.J., Robaire, B., and Trasler, J.M. (2007). Adverse Effects of 5-Aza-2′-Deoxycytidine on Spermatogenesis Include Reduced Sperm Function and Selective Inhibition of de Novo DNA Methylation. J Pharmacol Exp Ther 322, 1171–1180.

Oliva, M., Muñoz-Aguirre, M., Kim-Hellmuth, S., Wucher, V., Gewirtz, A.D.H., Cotter, D.J., Parsana, P., Kasela, S., Balliu, B., Viñuela, A., et al. (2020). The impact of sex on gene expression across human tissues. Science 369, eaba3066.

Peat, J.R., Dean, W., Clark, S.J., Krueger, F., Smallwood, S.A., Ficz, G., Kim, J.K., Marioni, J.C., Hore, T.A., and Reik, W. (2014). Genome-wide Bisulfite Sequencing in Zygotes Identifies Demethylation Targets and Maps the Contribution of TET3 Oxidation. Cell Reports 9, 1990–2000.

Rodriguez-Terrones, D., and Torres-Padilla, M.-E. (2018). Nimble and Ready to Mingle: Transposon Outbursts of Early Development. Trends Genet 34, 806–820.

Rulands, S., Lee, H.J., Clark, S.J., Angermueller, C., Smallwood, S.A., Krueger, F., Mohammed, H., Dean, W., Nichols, J., Rugg-Gunn, P., et al. (2018). Genome-Scale Oscillations in DNA Methylation during Exit from Pluripotency. Cell Syst 7, 63–76.e12.

Santi, D., Vincentis, S.D., Magnani, E., and Spaggiari, G. (2017). Impairment of sperm DNA methylation in male infertility: a meta-analytic study. Andrology-Us 5, 695–703.

Sherry, S.T., Ward, M.-H., Kholodov, M., Baker, J., Phan, L., Smigielski, E.M., and Sirotkin, K. (2001). dbSNP: the NCBI database of genetic variation. Nucleic Acids Res 29, 308–311.

Singmann, P., Shem-Tov, D., Wahl, S., Grallert, H., Fiorito, G., Shin, S.-Y., Schramm, K., Wolf, P., Kunze, S., Baran, Y., et al. (2015). Characterization of whole-genome autosomal differences of DNA methylation between men and women. Epigenet Chromatin 8, 43.

Smallwood, S.A., Lee, H.J., Angermueller, C., Krueger, F., Saadeh, H., Peat, J., Andrews, S.R., Stegle, O., Reik, W., and Kelsey, G. (2014a). Single-cell genome-wide bisulfite sequencing for assessing epigenetic heterogeneity. Nat Methods 11, 817–820.

Smallwood, S.A., Lee, H.J., Angermueller, C., Krueger, F., Saadeh, H., Peat, J., Andrews, S.R., Stegle, O., Reik, W., and Kelsey, G. (2014b). Single-cell genome-wide bisulfite sequencing for assessing epigenetic heterogeneity. Nat Methods 11, 817–820.

Smith, Z.D., Chan, M.M., Mikkelsen, T.S., Gu, H., Gnirke, A., Regev, A., and Meissner, A. (2012). A unique regulatory phase of DNA methylation in the early mammalian embryo. Nature 484, 339–344.

Souza, C.P.E. de, Andronescu, M., Masud, T., Kabeer, F., Biele, J., Laks, E., Lai, D., Ye, P., Brimhall, J., Wang, B., et al. (2020). Epiclomal: Probabilistic clustering of sparse single-cell DNA methylation data. Plos Comput Biol 16, e1008270.

Takashima, S., Takehashi, M., Lee, J., Chuma, S., Okano, M., Hata, K., Suetake, I., Nakatsuji, N., Miyoshi, H., Tajima, S., et al. (2009). Abnormal DNA Methyltransferase Expression in Mouse Germline Stem Cells Results in Spermatogenic Defects. Biol Reprod 81, 155–164.

Tang, W.W.C., Dietmann, S., Irie, N., Leitch, H.G., Floros, V.I., Bradshaw, C.R., Hackett, J.A., Chinnery, P.F., and Surani, M.A. (2015). A Unique Gene Regulatory Network Resets the Human Germline Epigenome for Development. Cell 161, 1453–1467.

Tucci, V., Isles, A.R., Kelsey, G., Ferguson-Smith, A.C., Group, the E.I., Tucci, V., Bartolomei, M.S., Benvenisty, N., Bourc’his, D., Charalambous, M., et al. (2019). Genomic Imprinting and Physiological Processes in Mammals. Cell 176, 952–965.

Uk, A., Collardeau-Frachon, S., Scanvion, Q., Michon, L., and Amar, E. (2018). Assisted Reproductive Technologies and imprinting disorders: Results of a study from a French congenital malformations registry. Eur J Med Genet 61, 518–523.

Vasiliauskaitė, L., Berrens, R.V., Ivanova, I., Carrieri, C., Reik, W., Enright, A.J., and O’Carroll, D. (2018). Defective germline reprogramming rewires the spermatogonial transcriptome. Nat Struct Mol Biol 25, 394–404.

Wen, L., and Tang, F. (2019). Human Germline Cell Development: from the Perspective of Single-Cell Sequencing. Mol Cell 76, 320–328.

WHO (2010). WHO laboratory manual for the examination and processing of human semen, FIFTH EDITION.

Wijchers, P.J., and Festenstein, R.J. (2011). Epigenetic regulation of autosomal gene expression by sex chromosomes. Trends Genet 27, 132–140.

Zhou, F., Wang, R., Yuan, P., Ren, Y., Mao, Y., Li, R., Lian, Y., Li, J., Wen, L., Yan, L., et al. (2019). Reconstituting the transcriptome and DNA methylome landscapes of human implantation. Nature 572, 660–664.

Zhu, P., Guo, H., Ren, Y., Hou, Y., Dong, J., Li, R., Lian, Y., Fan, X., Hu, B., Gao, Y., et al. (2018). Single-cell DNA methylome sequencing of human preimplantation embryos. Nat Genet 50, 12–19.

